# Riemannian Diffusion Kernel-smoothed Continuous Structural Connectivity On Cortical Surface

**DOI:** 10.1101/2025.09.08.674789

**Authors:** Lu Wang, Didong Li, Zhengwu Zhang

## Abstract

Atlas-free continuous structural connectivity has garnered increasing attention due to the limitations of atlas-based approaches, including the arbitrary selection of brain atlases and potential information loss. Typically, continuous structural connectivity is represented by a probability density function, with kernel density estimation as a common estimation method. However, constructing an appropriate kernel function on the cortical surface poses significant challenges. Current methods often inflate the cortical surface into a sphere and apply the spherical heat kernel, introducing distortions to density estimation. In this study, we propose a novel approach using the Riemannian diffusion kernel derived from the Laplace-Beltrami operator on the cortical surface to smooth streamline endpoints into a continuous density. Our method inherently accounts for the complex geometry of the cortical surface and exhibits computational efficiency, even with dense tractography datasets. Additionally, we investigate the number of streamlines or fiber tracts required to achieve a reliable continuous representation of structural connectivity. Through simulations and analyses of data from the Adolescent Brain Cognitive Development (ABCD) Study, we demonstrate the potential of the Riemannian diffusion kernel in enhancing the estimation and analysis of continuous structural connectivity.

## 1 Introduction

With the advancement of diffusion MRI (dMRI), we have started to uncover a vast amount of information about how different parts of the brain are connected with the dMRI-derived structural connectivity (SC). The SC quantifies how different brain regions are linked by networks of white matter fibers, which are essential for communication and information flow in the brain (Craddock et al., 2013; Fornito et al., 2013; Park & Friston, 2013). The most common way of representing the SC data is via a region-to-region graph model, where grey matter regions of interest (ROIs) are represented as nodes, and observed structural measurements serve as a proxy for edge strengths between nodes (Fornito et al., 2013). A critical aspect of this representation is the choice of ROIs, which typically relies on a predefined brain atlas parcellation (Allen et al., 2024; Arslan et al., 2018; Bryce et al., 2021).

However, this discrete, atlas-based framework for representing SC has notable drawbacks. First, the lack of consensus on a single optimal atlas-based representation leads to substantial variability. Multiple atlases exist (Desikan et al., 2006; Destrieux et al., 2010; Eickhoff et al., 2018; Glasser et al., 2016), and different parcellation schemes can yield inconsistent conclusions from the same underlying data (de Reus & Van den Heuvel, 2013; Zalesky et al., 2010). Second, aggregating sub-ROI-level information into coarse, ROI-to-ROI connectivity measures discards fine-grained detail (Consagra et al., 2024). Recent work (e.g., in Cole et al., 2021; Mansour L et al., 2022; Moyer et al., 2017) has sought to overcome these limitations by shifting from a discrete, ROI-based model to a continuous, parcellation-free representation. This continuous framework offers competitive reproducibility, improved predictive performance for various behavioral traits, and enhanced statistical power in detecting group differences in cognition, emotion, and sensory categories (Cole et al., 2021; Consagra et al., 2024; Mansour L et al., 2022).

The continuous SC model, first introduced by Moyer et al. (2017), aims to estimate a continuous intensity or density function that characterizes the distribution of streamline endpoints on the cortical surface. Unlike discrete models that assign connectivity masses to pairs of ROIs, the continuous density function defines a probability density over all pairs of points on the cortical surface, reflecting the likelihood that each pair is structurally connected via white matter fibers.

Existing approaches typically rely on kernel density estimation (KDE) to approximate this continuous density from streamline endpoint pairs, a task complicated by the cortical surface’s intricate geometry and the vast number of reconstructed streamlines. For instance, Moyer et al. (2017) and Cole et al. (2021) project the cortical surface onto a sphere and use a normalized spherical heat kernel for KDE. While the spherical geometry affords convenient closed-form solutions for heat kernels, mapping the complex cortical surface to a sphere inevitably introduces geometric distortions. In contrast, Mansour L et al. (2022) apply a squared exponential kernel with geodesic distances computed directly on the cortical surface mesh. Although this approach avoids spherical distortions, computing geodesic distances at high mesh resolutions is computationally burdensome (Li & Dunson, 2019; Tenenbaum et al., 2000).

Recent developments in manifold Gaussian processes facilitate direct KDE application on the native cortical surface manifold. Techniques introduced by Lindgren et al. (2011), and further developed by Borovitskiy et al. (2020) and Borovitskiy et al. (2021), allow the computation of Matérn kernels on compact Riemannian manifolds using the spectral theory of the Laplace-Beltrami operator (LBO). In a landmark study, Pang et al. (2023) demonstrated that leveraging the intrinsic geometry of the cortex, particularly through LBO eigenmodes, provides a more comprehensive characterization of brain dynamics than graph-based approximations. These geometric eigenmodes inherently account for cortical curvature and local spatial relations. Motivated by these advances, our work integrates white matter connectivity data with the cortical surface’s geometric information to achieve a more precise continuous representation of SC.

A crucial practical question in continuous SC analysis is determining how many streamlines are required to yield a reliable continuous representation. Generating an excessively large number of streamlines increases computational costs, hampers visualization, and adds storage demands. Yet, identifying an appropriate streamline count is nontrivial, as the needed number depends on multiple factors, including the downstream connectivity analysis tasks. While previous studies (Newlin et al., 2023; Roine et al., 2019) have examined the effects of streamline count on the reproducibility of graph-based measures in discrete SC frameworks, comparable guidelines for continuous SC models remain lacking. Given that continuous SC inherently smooths connectivity data, it may require fewer streamlines to achieve a stable and accurate representation.

In this study, we make two key contributions to the analysis of continuous SC. First, we propose an efficient method to directly estimate continuous SC density function on the complex cortical surface manifold. This is done through applying recently developed techniques for computing the Riemannian diffusion/Matérn kernel on compact Riemannian manifolds (Borovitskiy et al., 2020; Lindgren et al., 2011). We rigorously compare the continuous representations derived from the Riemannian diffusion kernel with those from the spherical heat kernel, evaluating their performance in terms of computational efficiency, connectome reproducibility, and predictive accuracy for cognitive traits. Second, we investigate the effect of streamline number on the quality of the estimated continuous SC. Our simulations demonstrate that the Riemannian diffusion kernel can infer a high-quality density function with fewer observations than the spherical heat kernel, showing the potential to reduce the computational burden of tractography without compromising connectivity’s estimation accuracy. Further analysis of data from the Adolescent Brain Cognitive Development (ABCD) Study reveals that the smoothed densities generated using the Riemannian diffusion/Matérn kernel produce lower prediction errors for oral reading ability, a crystallized cognitive trait, compared to those produced using the spherical heat kernel. Notably, the prediction errors remain stable even when the streamline count is reduced to approximately 50,000 per subject. Moreover, the significant local regions distinguishing between good and poor oral reading groups are preserved until streamline counts drop below this threshold, as assessed by a state-of-the-art statistical test of continuous SC for group differences.

Overall, our findings underscore the advantages of leveraging the cortical surface’s intrinsic geometry for continuous SC estimation, and provide practical recommendations for streamline reconstruction in the tractography step to ensure reliable and computationally efficient continuous SC modeling.

## 2 Methods

### 2.1 Dataset

In this study, we focus on a sample of 318 children from the ABCD Study (Casey et al., 2018), which aims to track brain development from childhood through adolescence in order to understand the biological and environmental factors that can affect the brain’s developmental trajectory. The participants in this study were aged 9 to 10 when they entered. The ABCD Study has released their neuroimaging data at two time points, spaced two years apart, which enables the assessment of both intra- and inter-individual variations in the mapped connectivity information.

In addition to the neuroimaging data, each subject also has a set of measurements related to cognitive traits. In our analysis, we focus on 4 age-adjusted scale scores from the NIH toolbox cognition domain: picture vocabulary test score, oral reading recognition test score, crystallized cognition composite score, and cognitive function composite score. The picture vocabulary test assesses participants’ ability to match words with corresponding photographic images presented on a computer screen using an audio recording of words. The reading recognition test evaluates participants’ ability to accurately read and pronounce letters and words. The crystallized cognition composite provides a global assessment of verbal reasoning and is derived by averaging the scores of the picture vocabulary and reading recognition tests. The cognitive function composite includes all the tests under the crystallized and fluid composites, providing a general measure of cognition for individuals. In our results, structural connectivity was less predictive of the fluid cognition composite than of the crystallized composite, regardless of whether smoothed or unsmoothed connectomes were used. Specifically, the correlation between predicted and observed scores was approximately 0.4 for the crystallized composite, compared to around 0.2 for the fluid composite. This pattern is also observed in Dhamala et al. (2021). Therefore, we omit the results for the fluid composite.

### 2.2 Imaging Data Preprocessing

For each subject, structural connectomes were constructed using both diffusion magnetic resonance imaging (dMRI) and T1-weighted MRI scans. The dMRI data capture the directionally dependent diffusion of water molecules, influenced by the underlying white matter microstructure, as water tends to diffuse more readily along neural fiber bundles (Bammer, 2003). By fitting smooth local models of these diffusion patterns, we identified white matter pathways through tractography (Cole et al., 2021). In parallel, T1-weighted images, which provide high contrast between gray and white matter tissues, were used to construct white matter surfaces and provide guidance on the growth of white matter fiber tracts.

All T1-weighted and dMRI datasets were obtained from the ABCD 4.0 release. Structural connectomes were generated using the surface-based connectome integration (SBCI) framework (Cole et al., 2021), a state-of-the-art dMRI preprocessing pipeline. SBCI constructs cortical surfaces and their spherical parameterizations via FreeSurfer (Fischl, 2012), representing each cortical hemisphere as a dense triangular mesh of 163,842 vertices. It then estimates white matter fiber tracts (or called streamlines in the literature) connecting the cortical surfaces using surface-enhanced tractography (St-Onge et al., 2018). On average, approximately 500,000 streamlines were generated per scan. Figure 1 shows the extracted streamlines and their endpoints on the cortical white surface for a randomly selected subject from the ABCD study.

**Figure 1.**
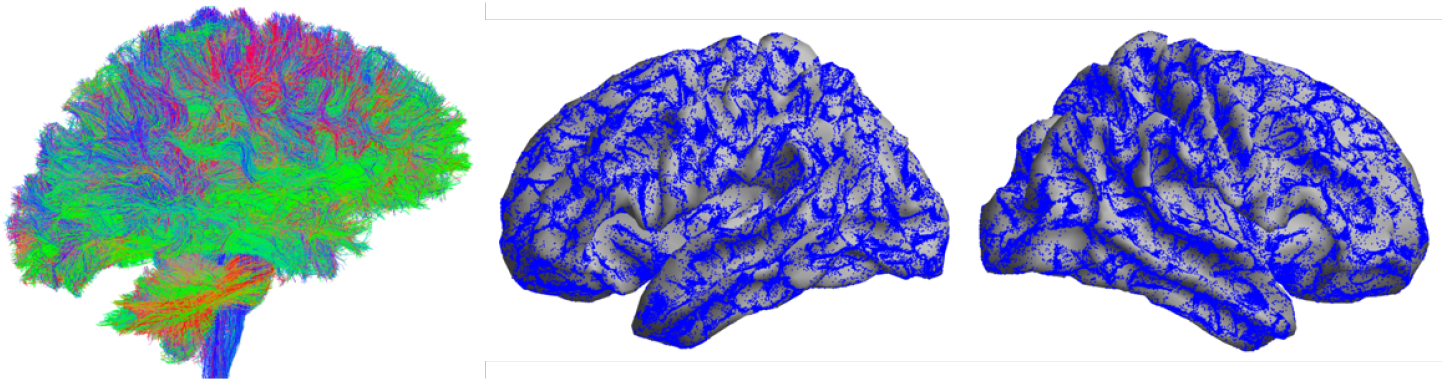
3D view of streamlines extracted from tractography and their endpoints on the cortical white surface for a randomly selected ABCD subject.

### 2.3 Continuous Structural Connectivity Estimation on Cortical Surface

Let Ω denote the white surface of the brain, representing the interface between cortical gray matter and white matter. In this work, we define *continuous structural connectivity* as a probability density function *p*(***x, y***) on Ω *×* Ω. For any two points ***x, y*** ∈ Ω, the function value *p*(***x, y***) specifies the probability density that ***x*** and ***y*** are connected by a white matter fiber. Since the streamlines are undirected, the order of the endpoints should not affect the density values; that is, the connection probability from ***x*** to ***y*** is the same as from ***y*** to ***x***, which gives the relation *p*(***x, y***) = *p*(***y, x***). Thus, *p* is a symmetric function mapping: Ω *×* Ω → [0, +∞).

For a single subject, let 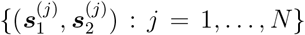 denote the set of observed streamline endpoint pairs on Ω *×* Ω. We treat these endpoint pairs as realizations drawn from the distribution governed by *p*(***x, y***). By symmetry, if 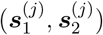 is observed, then its counterpart 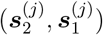 is also considered part of the distribution. Using KDE to estimate *p* based on these observations, we obtain the estimator

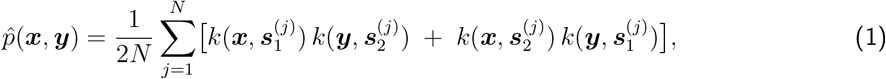

where *k*(·, ·) : Ω *×* Ω → [0, +∞) is a symmetric kernel function defined on the product space of the white surface. However, due to the complex geometry of Ω (as illustrated in Figure 1), the choices for *k*(·, ·) on Ω *×* Ω are limited.

#### 2.3.1 Related kernel functions

A series of recent works (Cole et al., 2021; Consagra et al., 2024; Moyer et al., 2017) addressed this challenge by mapping each hemisphere’s cortical surface to the unit sphere 𝕊 ^2^, thereby enabling KDE on the product space of unit spheres using the *spherical heat kernel* (SHK). The SHK generalizes the Gaussian kernel from Euclidean space to the unit sphere. For any ***u, v*** ∈ 𝕊^2^, the SHK is given by

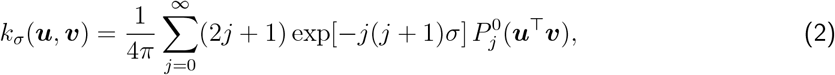

where *σ >* 0 is the bandwidth, and 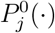 denotes the Legendre polynomial of degree *j* and order 0. The Legendre polynomials can be computed via the recurrence relation

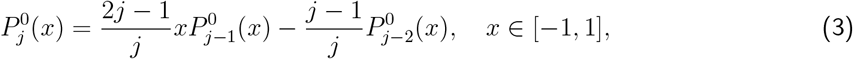

with initial conditions 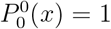 and 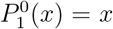. To numerically approximate (2), the infinite series is truncated after *J* terms. Because 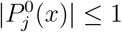 for *x* ∈ [−1, 1] and the coefficient (2*j* +1) exp[−*j*(*j* +1)*σ*] decreases as *j* grows for any *σ >* 0, we choose *J* such that (2*J* + 1) exp[−*J* (*J* + 1)*σ*] *<* 0.001. However, because the white surface and sphere have different intrinsic curvatures, this mapping cannot be strictly isometric and inevitably distorts distances between data points, which may affect downstrean analyses. Furthermore, although the mapping is designed to be a diffeomorphism, numerical approximations in its implementation can introduce additional errors, compounding the overall distortion.

An alternative approach that preserves aspects of the white surface geometry is to define *k*(·, ·) directly on Ω *×* Ω using a Gaussian function in geodesic distance (Mansour L et al., 2022):

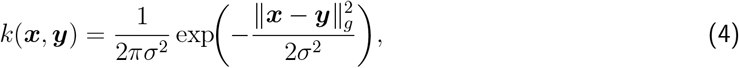

where ∥***x*** − ***y***∥_*g*_ is the geodesic distance along Ω. However, computing geodesic distances at scale is computationally expensive (Tenenbaum et al., 2000), and Feragen et al. (2015) and Da Costa et al. (2023) have shown that the geodesic squared exponential kernel (4) is not positive definite on nontrivial compact manifolds. In the next subsection, we introduce well-defined kernels on compact Riemannian manifolds that avoid these issues.

#### 2.3.2 Riemannian Matérn and diffusion kernels on the cortical surface

The Matérn family is a widely used class of covariance kernels for Gaussian processes (GPs) defined in Euclidean spaces. For ***x, x***′ ∈ ℝ^*d*^, a Matérn kernel (Borovitskiy et al., 2020) with parameters *σ*^2^, *κ, ν >* 0 is given by

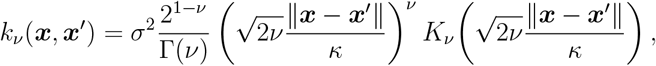

where *K*_*ν*_ is the modified Bessel function of the second kind. The parameter *σ*^2^ controls the GP variance, *κ* controls spatial correlation length, and *ν* controls smoothness (mean-square differentiability).

As *ν* → ∞, the Matérn kernel approaches the squared exponential kernel:

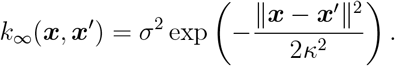

Consider now a compact, boundary-less Riemannian manifold ℳ of dimension *d* (e.g., a sphere or torus) with Riemannian metric *g*. Let *?*_*g*_ denote the Laplace-Beltrami operator (LBO) on ℳ. The Riemannian extension of the squared exponential kernel is often referred to as the heat or diffusion kernel. Throughout this paper, we use the term *Riemannian diffusion kernel* (RDK) for the squared exponential kernel defined on ℳ. Borovitskiy et al. (2020) proved the following result for the Riemannian Matérn and diffusion kernels for GPs.

##### Lemma 1

(Borovitskiy et al., 2020). *Let* {*λ*_*m*_} *be the eigenvalues of* −*?*_*g*_ *on* ℳ *and* {*f*_*m*_} *the corresponding eigenfunctions, so that*

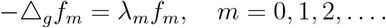

*Then the Riemannian Mat*é*rn and diffusion kernels on* ℳ *can be expressed as*

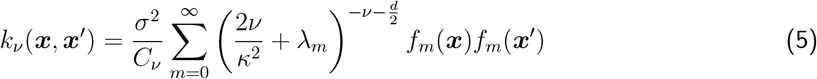

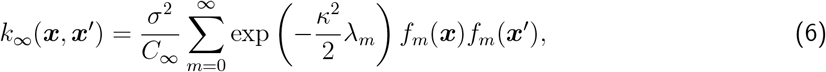

*where C*_*ν*_ *and C*_∞_ *are normalizing constants*.

Lemma 1 thus provides a spectral decomposition for the Riemannian Matérn and diffusion kernels, enabling pointwise evaluation on a discrete mesh approximation of a cortical surface.

The white surface Ω is represented as a union of two disjoint, compact, boundary-less Riemannian manifolds: Ω = Ω_*L*_∪Ω_*R*_, corresponding to the anatomically separate left and right brain cortical surfaces. This approach, which treats the surfaces as lacking a boundary, is a standard and practical simplification. While each hemisphere does have a physical boundary at the medial wall, treating the surfaces as closed allows us to use powerful and efficient numerical methods that are designed for boundary-less manifolds. This simplification is common in the field and is also used by methods that employ the spherical heat kernel after mapping the cortex to a sphere (Cole et al., 2021; Consagra et al., 2024; Moyer et al., 2017). In addition, to ensure connectivity estimates are zero within the medial wall region, we apply a standard medial wall mask to all estimated connectivity functions post-estimation. This guarantees that subsequent statistical analyses, visualizations, or interpretations are restricted to valid cortical grey matter regions, thereby avoiding contamination from artifacts associated with the medial wall boundary.

In practice, Ω_*L*_ and Ω_*R*_ are represented by fine triangular surface meshes. To compute the Riemannian Matérn/diffusion kernel matrix ***K***_*L*_ for vertices on the left hemisphere Ω_*L*_, we proceed as follows.

1. **Eigen-decomposition of the LBO:** Using the LaPy Python library (Reuter et al., 2006; Wachinger et al., 2015), we numerically approximate the eigenpairs {(*λ*_*m*_, *f*_*m*_)} of the LBO on Ω_*L*_:

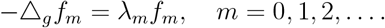

The eigenvalues satisfy 0 = *λ*_0_ *< λ*_1_ ≤ *λ*_2_ ≤, and the eigenfunctions {*f*_*m*_} form an orthonormal basis of the space of square-integrable functions on Ω_*L*_.
2. **Truncation and matrix construction:** To approximate the infinite sums in (5) or (6), we truncate after *M*_*T*_ terms. The number *M*_*T*_ can be chosen to equal the number of vertices in the surface mesh, which is the maximum number of LBO eigenpairs that can be computed numerically. Alternatively, *M*_*T*_ can be selected based on when the coefficient of the *M*_*T*_-th term in (5) or (6) approaches zero for a given bandwidth *κ*. Mostowsky et al. (2024) recommend using 1000 LBO eigenpairs by default to estimate the diffusion or Matérn kernel. In practice, we set *M*_*T*_ equal to the number of vertices in the surface mesh, which is *M*_*T*_ = 2562 throughout this paper. Let **Λ** be the diagonal matrix of the *M*_*T*_ smallest eigenvalues and ***F*** the matrix whose columns are the corresponding eigenfunctions evaluated at the vertices of mesh Ω_*L*_. For a suitable function Φ : ℝ → ℝ, we define Φ(**Λ**) by applying Φ to each diagonal entry of **Λ**. The kernel matrix on Ω_*L*_ then takes the form

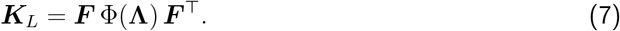

Using Lemma 1 with *d* = 2, choosing

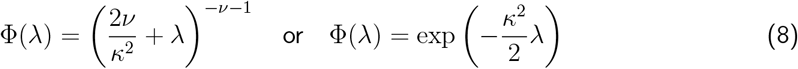

yields the Riemannian Matérn and diffusion kernel matrices, respectively.

Repeating the same procedure for the right hemisphere Ω_*R*_ yields ***K***_*R*_. If ***x*** and ***x***′ lie on different hemispheres, it is standard to set the kernel value *k*(***x, x***′) = 0 (Cole et al., 2021; Moyer et al., 2017).

Therefore, the kernel matrix for the entire white surface Ω is then

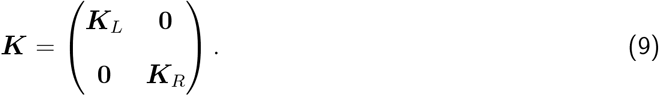

### 2.4 Bandwidth and Number of Streamlines

The bandwidth of a kernel function critically influences the results of KDE. In the RDK (6), the bandwidth parameter is *κ*, while in the SHK (2), the bandwidth parameter is *σ*. To investigate the effect of bandwidth, we consider a range of suitable values for each kernel. Specifically, we select a sequence of ten *κ*-values on a logarithmic scale from *κ*_min_ = 0.6252 to *κ*_max_ = 18.9204, and a sequence of ten *σ*-values from *σ*_min_ = 0.0001 to *σ*_max_ = 0.05. These sequences are chosen such that the effective “influence radii” of the two kernels, when viewed on a sphere, are comparable. Figure 2 illustrates the kernel values from a fixed vertex to all other vertices on the left hemisphere for both the RDK and the SHK, under varying bandwidths.

**Figure 2.**
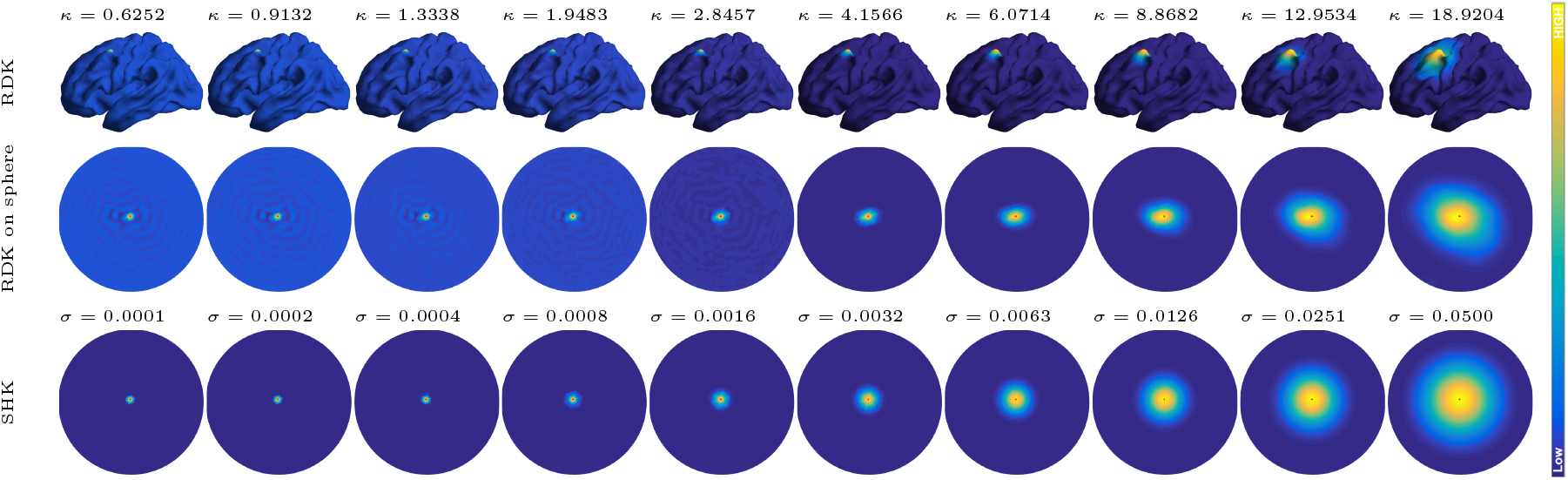
The Riemannian diffusion kernel (RDK) *k*_*κ*_(***v***, ·) defined on the white surface (top row) and projected onto a sphere (middle row) for different *κ*-values, and the spherical heat kernel (SHK) *k*_*σ*_(***v***, ·) defined on 𝕊^2^ (bottom row) for various *σ*-values. The vertex ***v*** is indicated by a red dot, and colors represent the kernel values.

Another practical factor affecting *p*(***x, y***) estimation is the sample size, i.e., the number of streamlines connecting cortical surface in tractography. Previous studies have suggested that millions of streamlines may be necessary to confidently recover structural connectivity (Smith et al., 2015; St-Onge et al., 2018). The number of streamlines is influenced by the seeding procedure: increasing seeds yields more observed streamlines, enhancing the reliability of microstructural estimates but at a higher computational cost for generation, visualization, and storage.

To determine how many streamlines are needed for stable and reliable continuous structural connectivity estimates, we vary the number of streamlines and assess the resulting changes in both prediction and inference tasks. With the ABCD dataset described in Section 2.1, each subject’s tractography initially included, on average, about 5.2 *×* 10^5^ streamlines (SD ≈ 5.6 *×* 10^4^). This magnitude is consistent with prior work (Cole et al., 2021) focusing on continuous structural connectivity and provides a good balance between computational efficiency and anatomical coverage. We then downsampled the streamlines for each subject to 50%, 25%, 10%, 5%, and 1% of the original count. We use a series of criteria, including the trait prediction and local inference test described in the next subsections, to determine the adequate number of streamlines to construct.

### 2.5 Evaluation Metrics

Although various criteria have been proposed for parameter tuning (e.g., bandwidth selection) in KDE, there is no universal consensus. In this work, tailored to our connectivity applications, we compare four criteria to evaluate the estimated brain connectivity: the distance-based intraclass correlation coefficient (dICC), reliability *µ*_*intra*_, identifiability (Mansour L et al., 2022) and cognitive trait prediction errors. Each of these criteria emphasizes different aspects of the kernel smoothing process.

#### 2.5.1 Distance-based intraclass correlation coefficient (dICC)

The dICC (Xu et al., 2021) measures the reliability of complex data objects, such as functions or matrices. It extends the classical intraclass correlation coefficient (ICC), originally defined for scalar observations, to more general settings.

Suppose we have *n* subjects, each measured *n*_*i*_ times, and let *X*_*ij*_ denote the *j*-th observation from subject *i*. Define:

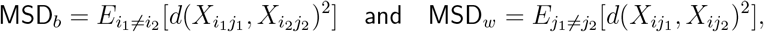

where *d*(·, ·) is a distance measure. The dICC is then computed as

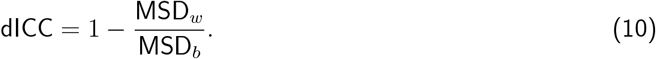

In practice, we estimate MSD_*b*_ and MSD_*w*_ using sample averages:

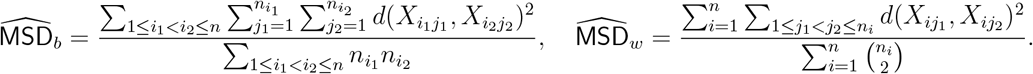

For the ABCD dataset, we have 318 children scanned at two time points (separated by two years), yielding both intra- and inter-individual variation. We assess reliability by applying the dICC to the continuous connectivity estimates derived at different bandwidths. The distance *d*(·, ·) between two discretized high-resolution connectivity matrices is computed using the Frobenius norm. A dICC value below 0.4 suggests poor reliability.

#### 2.5.2 Reliability *µ*_*intra*_ and identifiability

Mansour L et al. (2022) proposed two metrics for evaluating kernel bandwidth: *reliability µ*_*intra*_ and *identifiability*. Reliability *µ*_*intra*_ is the mean intra-individual similarity, reflecting the consistency of an individual’s connectome across repeated measurements. Identifiability assesses the extent to which individuals can be distinguished from one another. It is defined as:

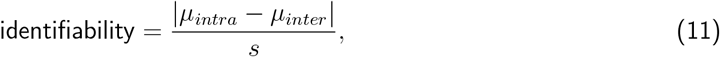

where *µ*_*inter*_ is the mean inter-individual similarity and *s* is the pooled standard deviation of similarities (Heumann et al., 2022). Similarities are computed as the average of row-wise Pearson correlations between connectivity matrices. High reliability and identifiability indicate that smoothing with the chosen bandwidth preserves individual differences while maintaining within-subject consistency.

#### 2.5.3 Trait prediction

Another approach is to select the bandwidth that optimizes prediction accuracy for downstream tasks, such as cognitive trait prediction. For the ABCD dataset, we use the baseline year continuous structural connectivity to predict cognitive measurements of 318 subjects. Each subject’s continuous connectivity is a high-dimensional object, and to reduce its dimensionality, we apply principal component analysis (PCA), extracting up to 317 principal components (PCs).

We then fit a ridge regression model to predict each cognitive trait from the first *K* PCs, where *K* ranges from 10 to 317 (in increments of 10). The ridge penalty is tuned via cross-validation. For each trait, we conduct 50 random 80-20 train-test splits, record the mean squared error (MSE) and prediction–outcome correlation on the test sets, and select the bandwidth that yields the best out-of-sample performance (lowest MSE or highest correlation).

### 2.6 Identifying Key Regions through Local Inference Testing

Beyond overall predictive performance, we may also want to identify where cognitive differences manifest locally in the brain’s connectivity structure. To this end, we apply a state-of-the-art local inference test proposed by Consagra et al. (2024). This test detects regions in Ω *×* Ω where continuous connectivity differs between two groups (e.g., individuals with high vs. low cognitive ability).

First, approximate each subject’s continuous connectivity function *p*_*l*_ (for group *l* = 1, 2) using a reduced-rank representation:

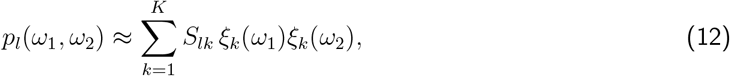

where {*ξ*_*k*_} are orthonormal basis functions with local support on Ω, and {*S*_*lk*_} are low-dimensional coordinates representing each subject’s connectivity in a *K*-dimensional space. Consagra et al. (2024) showed that testing for differences in *S*_1*k*_ vs. *S*_2*k*_ (for *k* = 1, …, *K*) can pinpoint the areas of Ω *×* Ω where group differences occur. By performing *K* univariate tests on the coefficients and using a correction for multiple comparisons (Holm, 1979), one can identify a region 𝒞 ⊆ Ω *×* Ω where connectivity differs between the two groups with controlled error rates.

This local inference procedure complements the other evaluation metrics by revealing not just how well we can predict cognitive traits or measure reliability, but also where, spatially, these differences arise within the brain’s connectivity landscape.

### 2.7 Other Implementation Details

#### 2.7.1 Cortical surface representation

Any computation or storage of the continuous connectivity 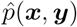 in (1) requires some form of discretization. Current approaches discretize the continuous connectome by forming pointwise estimates over all pairs of vertices in a high-resolution mesh grid on Ω (Consagra et al., 2024). To manage data complexity and computational resources, we use a standard downsampled fsaverage-ico4 surface mesh (Feilong et al., 2024) from the dense FreeSurfer fine mesh, comprising 2,562 vertices per hemisphere. Although higher-resolution meshes, such as fsaverage-ico6 with 40,962 vertices per hemisphere, could provide more detailed representations, the chosen resolution offers a practical balance between spatial detail and resource demands. For example, storing a single 2, 562 *×* 2, 562 connectivity matrix requires about 50 MB, while a 40, 962 *×* 40, 962 matrix would require approximately 12.5 GB. Connectomes exceeding several gigabytes pose significant challenges for data handling and downstream analyses, including extended loading times and substantial computational overhead. Even with high-performance computing capabilities, the marginal benefits of extremely high-resolution connectomes may not justify their considerable resource costs (Mansour L et al., 2022).

The continuous structural connectivity sampled on a mesh grid with *M* vertices can be represented by a large *M × M* connectivity matrix ***P***, where each element is a pointwise density estimate from (1) for a pair of vertices. The computation of ***P*** using a convientional KDE method poses a challenge due to its computational complexity of *O*(*M* ^2^*N*), particularly given that *M* usually ranges in the thousands and *N* in the millions. Moreover, this computation needs to be performed separately for each individual scan, based on their precise streamline endpoint data. This results in a substantial computational burden, necessitating the development of efficient algorithms to address the challenge.

#### 2.7.2 Fast estimation of continuous connectivity

We employ the efficient smoothing algorithm from Mansour L et al. (2022) to compute the large connectivity matrix ***P***, reducing the computational complexity from *O*(*M* ^2^*N*) to *O*(*M* ^3^), where the number of vertices *M* on a mesh grid is typically much smaller than the number of streamlines *N*. The algorithm first maps the streamline endpoints to the nearest vertices of the surface mesh, striking a compromise between precision and computational complexity. Following this, we can pre-store the values of *k*(·, ·) at each pair of vertices in an *M × M* matrix ***K***. Additionally, we introduce two half-incidence matrices ***U*** and ***V***, each of size *M × N*, to encode the streamline endpoint information. Specifically, if the *k*th streamline ends at vertices ***v***_*i*_ and ***v***_*j*_, then the *k*th columns of ***U*** and ***V*** consist of vectors with a single nonzero element 1 located at ***U*** (*i, k*) and ***V*** (*j, k*) respectively. Mathematically, the large connectivity matrix ***P*** can be computed by

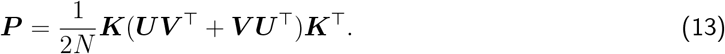

Define

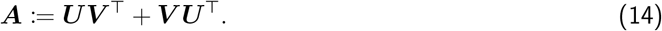

Note that ***A*** is actually a symmetric *M × M* adjacency matrix, with each entry ***A***(*i, j*) storing the streamline count between nodes ***v***_*i*_ and ***v***_*j*_. Plugging in (14) into (13), we have

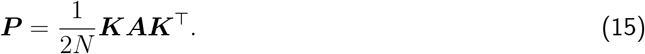

The scheme (15) eliminates the need for nested for-loops by utilizing formula (1), achieving a computational complexity of only a product of three *M × M* matrices. When smoothing multiple scans, the kernel matrix ***K*** is computed just once at the beginning of the procedure, since all subjects share the same surface mesh after registration alignment. In addition, the cost of deriving the discrete adjacency matrix ***A*** from the tractography of an individual scan is practically negligible. This leads to significant computational savings compared to the accurate approach (1) based on precise streamline endpoint locations.

## 3 Results

### 3.1 Evaluating the Estimated Laplace-Beltrami Eigenpairs on White Surface

To compute the Riemannian diffusion/Matérn kernels, the first step is to obtain LBO eigenvalue and eigenfunction pairs. We first validate the accuracy of the estimated eigenvalues of the LBO on the cortical white surface using Weyl’s law (Zelditch, 2017).

#### Lemma 2

(Weyl’s law). *Let* (ℳ, *g*) *be a compact Riemannian manifold of dimension d, with Laplace eigenvalues* 0 = *λ*_0_ *< λ*_1_ ≤ · · ·. *Then*

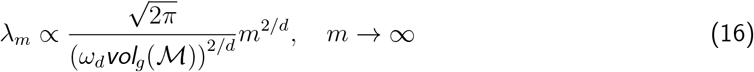

*where ω*_*d*_ *is the volume of the unit ball in* ℝ^*d*^, *and vol*_*g*_(ℳ) *is the Riemannian volumn of* ℳ *(e*.*g*., *surface area for d* = 2).

As the intrinsic dimension of the white surface on each hemisphere is *d* = 2, Lemma 2 indicates that its LBO eigenvalues *λ*_*m*_ asymptotically follow the linear scaling *λ*_*m*_ ∝ *m*. Equivalently, under logarithmic transformation, log(*λ*_*m*_) grows linearly in log(*m*) with a slope of 1. Figure 3 displays the log-log plot of the first 2562 eigenvalues of LBO on Ω_*L*_ estimated on fsaverage-ico6 mesh and fsaverage-ico4 mesh respectively. The fsaverage-ico6 mesh, with 40962 vertices on each hemisphere, more accurately approximates the white surface compared to the lower resolution fsaverage-ico4 mesh, which has 2562 vertices on each hemisphere. The plot shows that the first 2562 LBO eigenvalues estimated on fsaverage-ico6 mesh follow Weyl’s law closely, while the latter eigenvalues estimated on fsaverage-ico4 mesh grow much faster than a linear rate.

**Figure 3.**
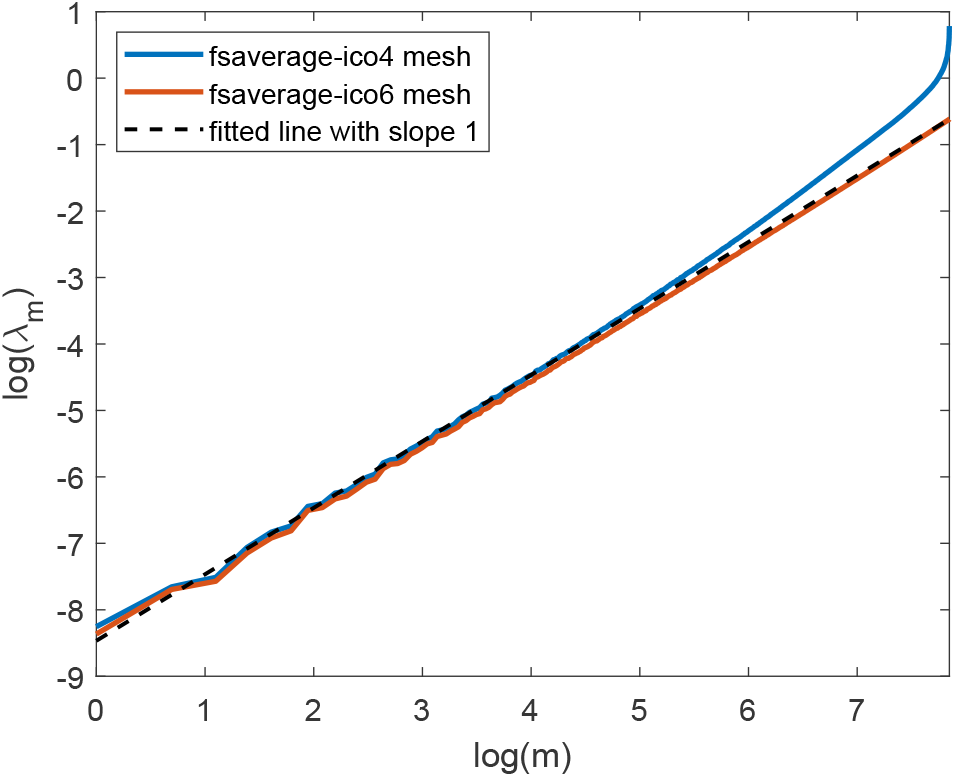
Logarithms of the first 2562 eigenvalues of the Laplace-Beltrami operator on Ω_*L*_ estimated using fsaverage-ico4 mesh (2562 vertices) and fsaverage-ico6 mesh (40962 vertices).

The eigenfunctions estimated on fsaverage-ico6 mesh also appear more reliable than those on the fsaverage-ico4 mesh as displayed in Figure 4. The figure indicates that the latter eigenfunctions estimated on the fsaverage-ico4 mesh tend to approach zero over a larger portion of the surface. Previously, we chose the fsaverage-ico4 mesh for efficiency in estimating continuous structural connectivity, as discussed in Section 2.7.1. To leverage more accurate results from the higher resolution fsaverage-ico6 mesh, we downsampled the first 2562 eigenvectors computed on this mesh to the fsaverage-ico4 mesh using the mapping between the two meshes. These downsampled eigenvectors serve as approximations to the first 2562 eigenfunctions of the LBO on each unihemispheric white surface. These eigenvectors form the matrix ***F*** in Equation (7).

**Figure 4.**
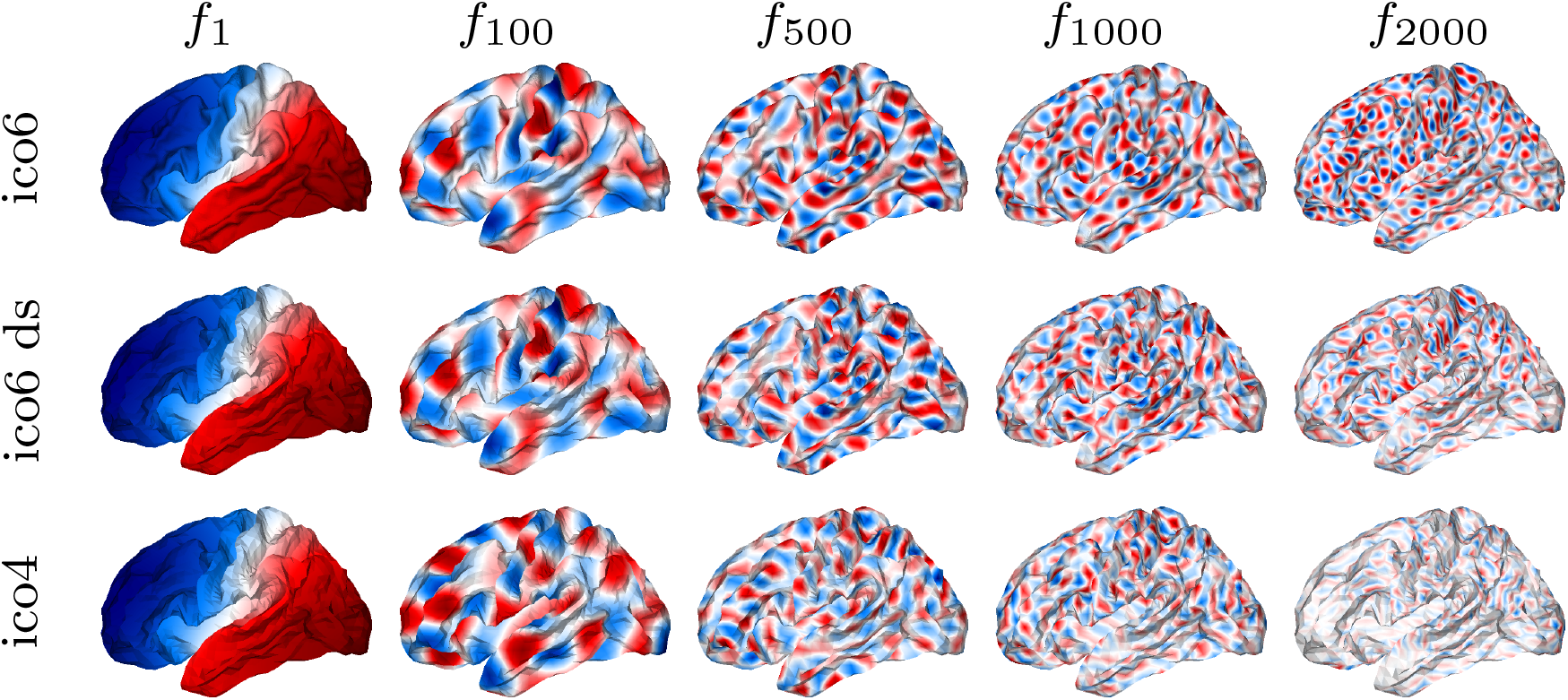
Eigenfunctions *f*_1_, *f*_100_, *f*_500_, *f*_1000_, *f*_2000_ of the Laplace-Beltrami operator on Ω_*L*_ estimated using fsaverage-ico6 mesh (upper), ico6-downsample-to-ico4 mesh (middle), and fsaverage-ico4 mesh (bottom). The value of the eigenfunction is given by the color.

### 3.2 Computing Efficiency

After obtaining the estimated eigenvalues and discretized eigenfunctions of the LBO of the white surface, the computation of the RDK matrices, ***K***_*L*_ and ***K***_*R*_, is more time-efficient than the computation of the SHK matrices. This is because computing the RDK matrix for each bandwidth *κ* involves simply multiplying three matrices, as detailed in (7). In contrast, there are two main reasons why the computation of the SHK matrices is slower. Firstly, the number of components in the finite truncation of SHK in (2) must be very large for small bandwidth values of *σ* to ensure that the coefficient (2*J* +1) exp[−*J* (*J* +1)*σ*] falls below the small threshold of 0.001 at truncation *J*. For example, *J* = 368 at the smallest value of *σ* = 0.0001 and *J* = 14 at the largest value of *σ* = 0.05. Secondly, the evaluation of the Legendre polynomial functions 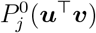 for each pair of vertices in SHK (2) is time-consuming, as 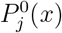 is computed iteratively according to (3).

Figure 5 displays the average CPU time over 10 replications for computing the high resolution (2562 *×* 2562) kernel matrices ***K***_*L*_ and ***K***_*R*_ in (9), for both RDK and SHK across their respective 10 bandwidths as specified in Section 2.4. These experiments were conducted using Matlab 2024a on a single CPU. For RDK, we only pre-stored the eigenvalues and the eigenvector matrix ***F*** in (7). For SHK, we precomputed the pairwise dot products ***u***^⊤^***v*** for each pair of vertices on the spherical mesh before conducting the experiments. The figure indicates that the computation of RDK matrices takes less than 1 second across all the bandwidths, whereas the computation of SHK matrices takes more than 10 seconds at very small bandwidth.

**Figure 5.**
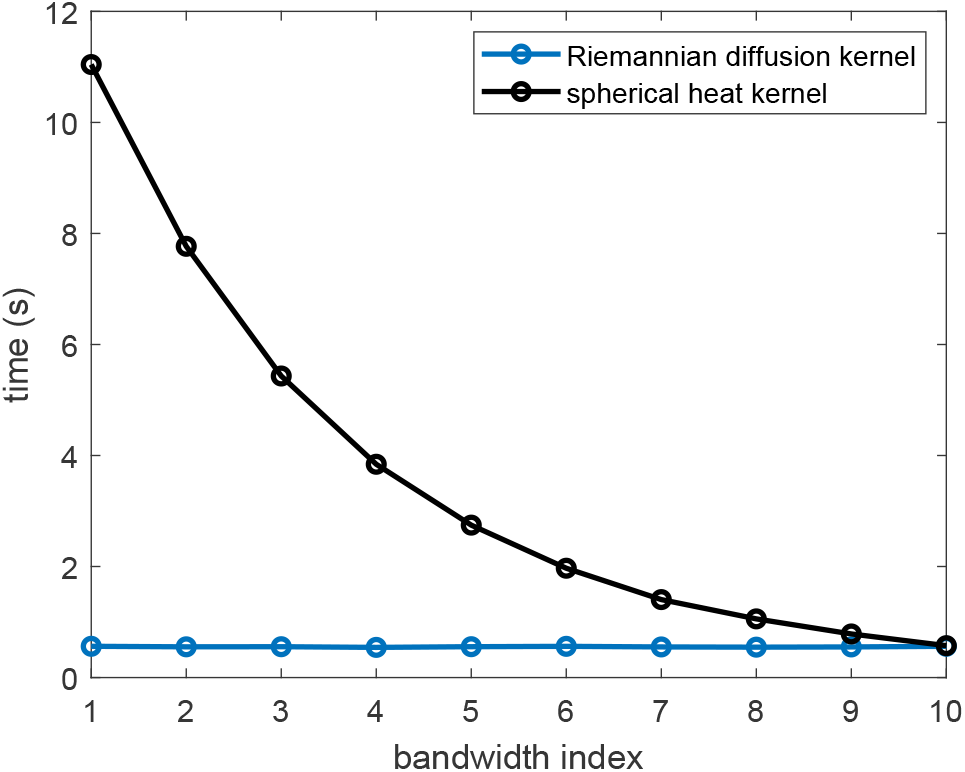
Average CPU time over 10 replications for computing the kernel matrices ***K***_*L*_ and ***K***_*R*_ in (9) for two kernels across 10 pre-specified bandwidths.

Regarding the computation of the SHK matrices, we found that recomputing the Legendre polynomial values for each bandwidth is actually faster than precomputing and storing these values for every pair of vertices on the unit sphere mesh. This is primarily due to the substantial memory overhead of caching, which scales as *O*(*M* ^2^*J*_max_), where *M* is the number of mesh vertices and *J*_max_ is the truncation index corresponding to the smallest bandwidth. Specifically, caching the polynomials 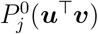 up to *J*_max_ = 368 for all vertex pairs consumed approximately 37.55 GB of memory, compared to just 1.86 GB when computing kernel values directly without caching. Because the caching approach involves allocating and processing a massive multi-gigabyte matrix – far too large to fit in the CPU’s fast cache – the processor spends most of its time waiting for data to be transferred from the much slower system RAM. Consequently, the speed bottleneck is memory access rather than computation. In practice, the massive memory overhead of caching negates its theoretical benefits, resulting in slower performance compared to the more memory-efficient direct computation approach. For example, preparing the cache alone took an average of 42 seconds, while computing the kernel matrix from scratch at the smallest bandwidth took less than 12 seconds. Moreover, Figure S1 in the supplementary material shows that the cached approach remains slower than the direct method across all bandwidths.

After obtaining the bi-hemispherical kernel matrix ***K*** as described in (9) for each kernel, the kernel-smoothed connectivity matrix ***P*** can be subsequently computed via (15). While the theoretical computational complexities of ***P*** for both RDK and SHK are identical, practical applications reveal that the computation of ***P*** is significantly more efficient when using RDK compared to SHK. This discrepancy is shown in Supplementary Figure S2, which illustrates that the average CPU time required to compute the RDK-smoothed ***P*** for each scan is consistently shorter than that for the SHK-smoothed connectivity matrix across all bandwidths. As a result, computing the RDK-smoothed connectivity matrices for 318 ABCD subjects across the 10 pre-specified bandwidths saves approximately 1 hour and 10 minutes of CPU time compared to computing the corresponding matrices using SHK.

### 3.3 Simulation Study: Evaluating Consistency in Kernel Density Estimation

In this section, we assess the consistency of each kernel method by checking whether the error between the true and estimated densities decreases consistently as the number of observed streamlines increases. To accomplish this, we generate varying numbers of streamline endpoints from a known two-bundle mixture distribution on the white surface. Detailed information regarding the design and results of the simulation study can be found in Supplementary Section S1. The results indicate that the estimation error consistently decreases with increasing sample sizes for each kernel, confirming their consistency. Additionally, for this simple mixture distribution where streamlines form two bundles, RDK achieves the lowest average estimation error when the sample size is small, specifically when there are fewer than 200 streamlines in total or fewer than 100 streamlines per bundle. The Riemannian diffusion/Matérn kernels also exhibit significantly better time efficiency than SHK in computing kernel-smoothed connectivity matrices for each specified number of streamlines.

Figure 6 provides a direct visualization of how well the estimated densities recover the true density. We randomly generate 100 streamline endpoint pairs from the true density, and display the estimated density on the white surface for both RDK and SHK. Compared to the true density shown in panel (a) of Figure 6, RDK provides a more accurate estimation, while SHK exhibits distinct errors on the neighboring gyrus around the high-density region at the sulcus. Panel (d) shows that the SHK method underestimates the density values for many vertices.

**Figure 6.**
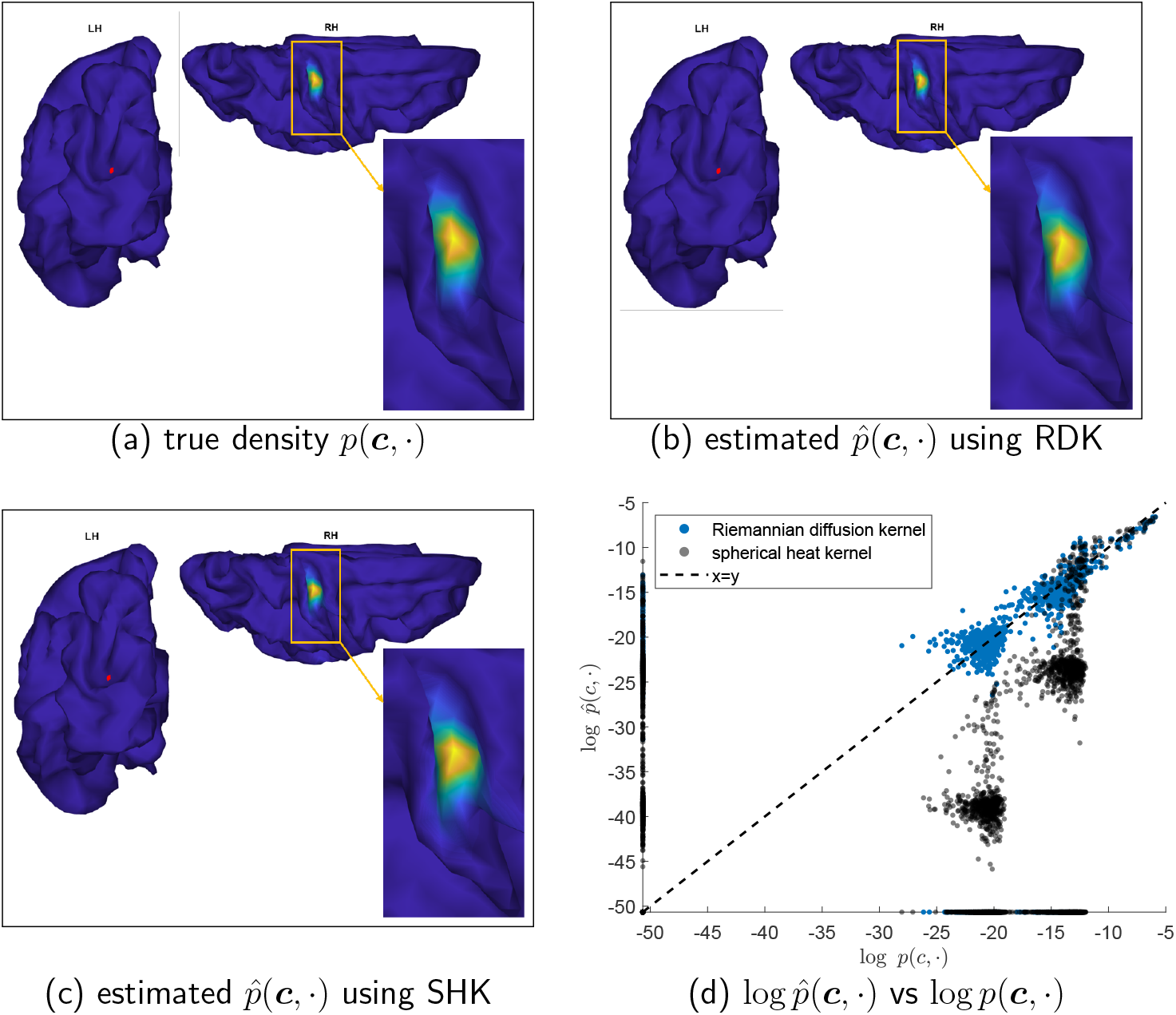
Comparison of the true density function, *p*(***c***, ·), with estimated densities on the white surface in panels (a)-(c) using a dataset generated with 100 streamlines. The point ***c*** is denoted by a red dot, and the color represents the function value. For clarity, a high density region is highlighted in orange in each panel, and a zoomed-in view of this region is presented in the bottom right corner. Panel (d) shows the estimated density values plotted against the true values on a logarithmic scale for each vertex of the surface mesh, using different kernels.

### 3.4 Bandwidth Selection

Using the streamline endpoint data observed from 318 subjects in the ABCD dataset at two different years, we compared the performances of the four criteria discussed in Section 2.5 for selecting the bandwidth in estimating continuous connectivity. In the following context, the high-resolution connectome without smoothing refers to the adjacency matrix based on the streamline count defined in (14).

#### 3.4.1 The dICC measure

Figure 7 displays the dICC values for the RDK and SHK estimated continuous connectivity under 10 bandwidths, respectively. We found that kernel smoothing improves reliability. In contrast, high-resolution connectomes without smoothing exhibited a relatively low reliability (dICC *<* 0.5). Notably, for both kernels, intermediate kernel bandwidths (e.g., the 4th, 5th, and 6th) led to a substantial increase in reliability, while excessively small or large bandwidths only resulted in marginal increases in dICC.

**Figure 7.**
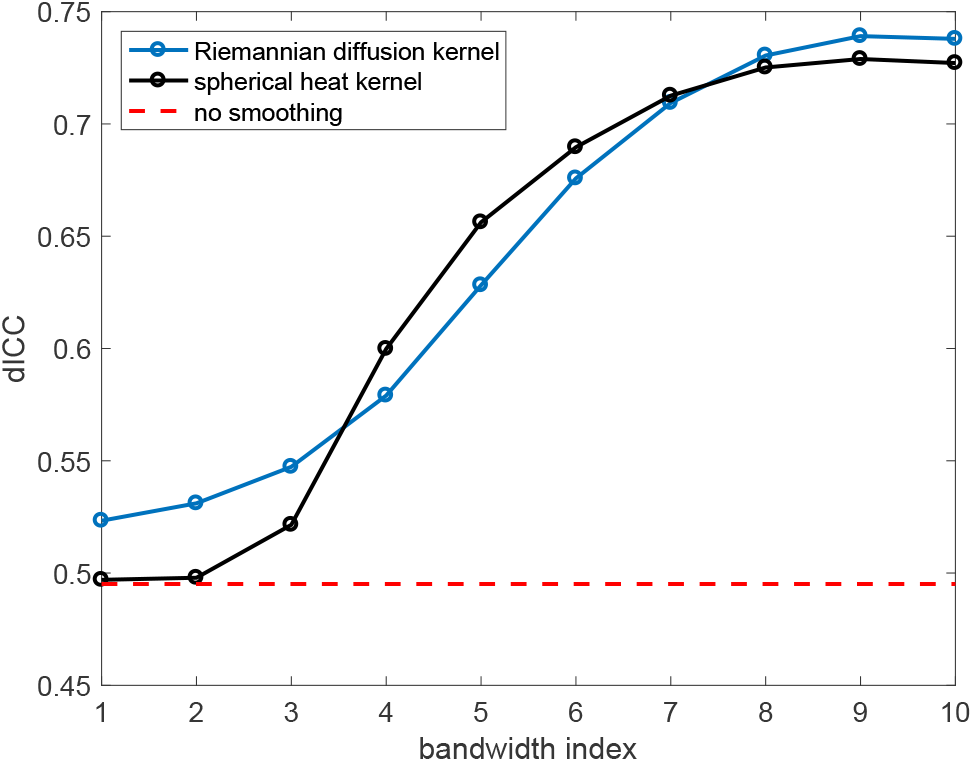
The dICC (reliability measure) values for kernel smoothed connectivity matrices (15) using two different kernels and 10 bandwidths for each.

In our study, a dICC *>* 0.5 implies that the mean squared differences for continuous connectivity representations between subjects are greater than those within subjects. However, considering the young age of the subjects in the ABCD study and their ongoing brain development, the differences in structural connectomes over a two-year interval may be significant for an adolescent. Therefore, striving for the highest dICC value might not be reasonable. We conducted visual inspections in Figure 8 to assess the actual smoothing impacts of different bandwidths. Figure 8 shows that the largest bandwidth value for either kernel results in excessive smoothing and severe distortion of signal peaks, despite achieving almost the highest dICC for kernel smoothed connectivity matrices. Therefore, we cannot rely solely on the dICC measure to select the optimal bandwidth.

**Figure 8.**
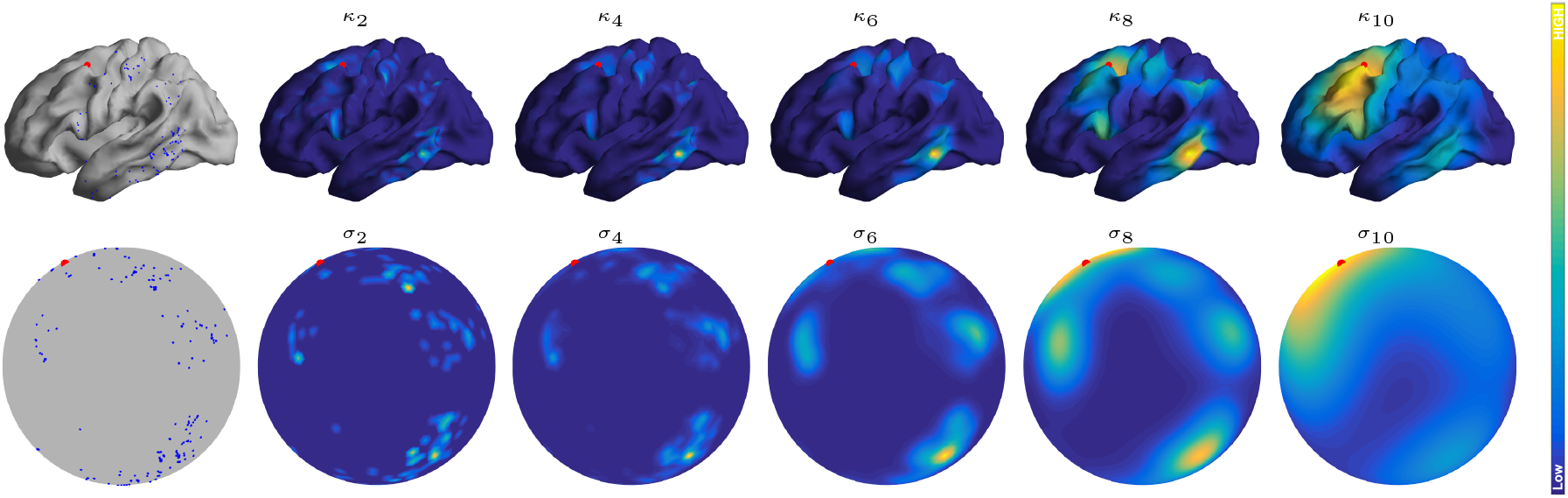
Visualization of the estimated connectivity density function 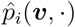 of a randomly selected subject, on the white surface using the Riemannian diffusion kernel (first row) and on a sphere using the spherical heat kernel (second row) over different bandwidths. The color gives the function value, with the vertex ***v*** marked with a large red dot. The first column displays the endpoint locations (blue dots) of the streamlines connected to the vertex ***v***.

#### 3.4.2 Reliability *µ*_*intra*_ and identifiability

Figure 9 summarizes the impact of bandwidth on the reliability *µ*_*intra*_ and identifiability (11) of high-resolution connectomes. We observed that increasing the bandwidth consistently improves the reliability *µ*_*intra*_ (mean intra-subject similarity), but simultaneously decreases the identifiability. This suggests that an excessively large bandwidth not only reduces the discrepancies between the connectomes observed at different ages within the same adolescent, but also diminishes inter-individual differences due to the overly smoothing effect illustrated in Figure 8. Consequently, this results in a more uniform connectome population, which is not desirable.

**Figure 9.**
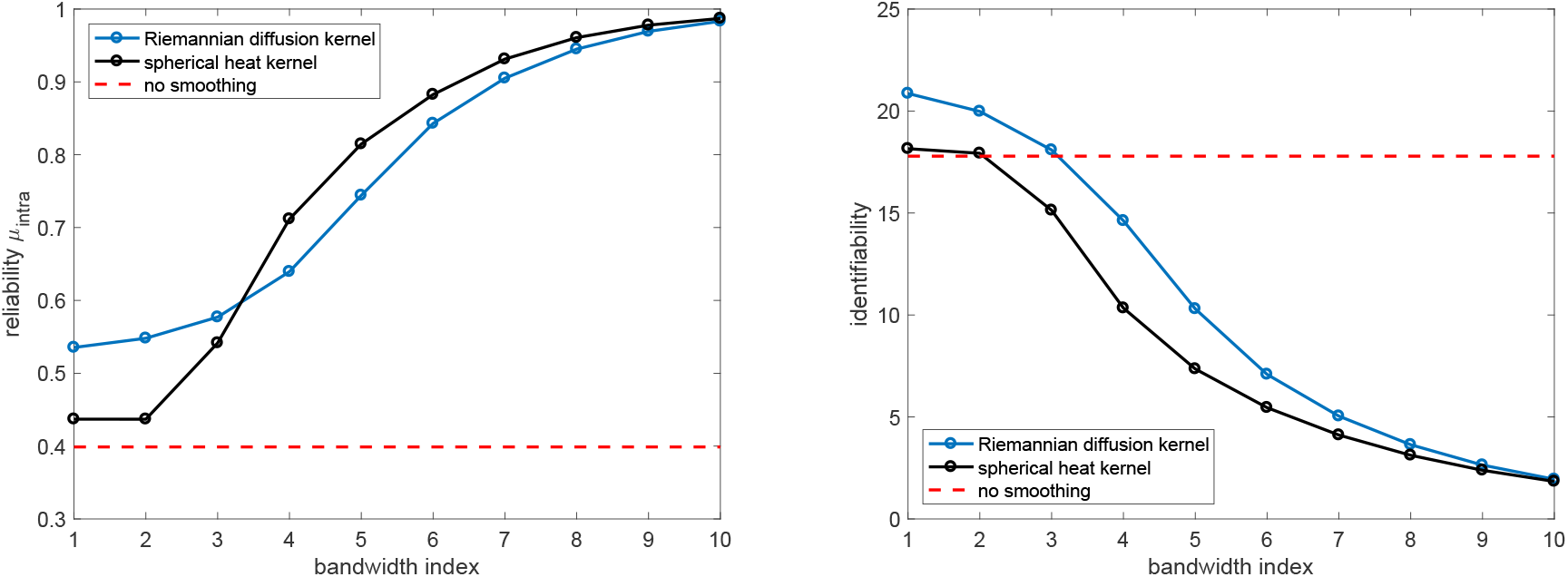
The reliability *µ*_*intra*_ (left) and identifiability (right) values for kernel smoothed connectomes using two different kernels across a set of 10 bandwidths.

Moreover, while higher reliability generally indicates greater similarity between a subject’s connectomes across two time points, this may not be ideal for the ABCD dataset. Given the two-year interval between scans, subtle neurodevelopmental changes are expected. If the SHK method obscures these fine-grained differences, causing the two connectomes to appear more similar than they truly are, this could artificially inflate within-subject similarity. Thus, maximizing reliability alone can be counterproductive, as it may come at the cost of erasing meaningful developmental signals. Therefore, selecting the optimal bandwidth requires a careful balance between reliability *µ*_*intra*_, identifiability, and qualitative assessment through visual inspection.

#### 3.4.3 Cognitive trait prediction

In this section, we compare the performance of different kernel methods for predicting cognitive traits from the smoothed connectomes. This approach has the potential to address the limitations of earlier criteria, and hence we additionally present results from using the Riemannian Matérn kernels (RMKs). As introduced in Section 2.3.2, RMKs can also be applied to Riemannian manifold. The key distinction between the density estimates provided by RMKs and RDK lies in their differentiability: RDK yields density functions that are infinitely differentiable, while those estimated using RMKs are finitely differentiable. This finite differentiability can be fine-tuned by adjusting the smoothness parameter *ν* in (5), thereby offering greater flexibility in certain scenarios.

Figure 10 displays the average out-of-sample MSE for predicting the four cognitive measurements listed in Section 2.1, across different bandwidths for each kernel method (see Supplementary Figure S6 for the correlation results). We observe that all the kernel smoothing methods achieve better predictive performance at an optimal bandwidth than without smoothing in predicting the four cognitive traits, although the optimal bandwidth varies depending on the specific cognitive trait being predicted. Additionally, Figure 10 shows that as the smoothness parameter *ν* increases, the behavior of RMK approaches that of RDK.

**Figure 10.**
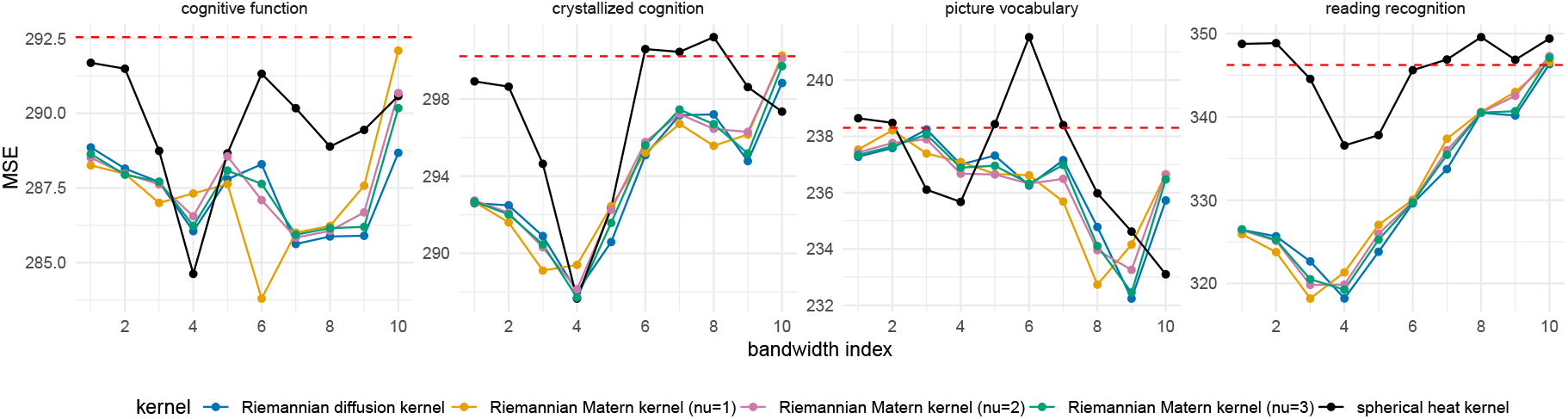
Average out-of-sample MSE between the observed and predicted cognitive measures, based on 50 random train-test splits, for five kernel methods across 10 bandwidths. The red dashed line represents the results for connectomes with no smoothing.

In predicting reading recognition scores, RDK and RMKs clearly outperform SHK based on both out-of-sample MSE and correlation. Furthermore, all the kernels achieved the lowest MSE and the highest correlation on average from the test data at the fourth bandwidth grid point, except for RMK with *ν* = 1, which achieved this at the third bandwidth point.

For picture vocabulary scores, all the kernels appeared to achieve the best performance at very large bandwidth values, with SHK performing best at the largest bandwidth. Visual inspection in Figure 8 revealed that the kernel-smoothed connectome at extremely large bandwidths suffers from over-blurring issues, which could lead to unreliable prediction results. Regarding the crystallized cognition composite and the cognitive function composite (total composite), each kernel’s predictive curves over bandwidths exhibited similar patterns to those of reading recognition and picture vocabulary. This was expected due to the relationship between core cognition measures and the composite scores.

Table 1 reports the average out-of-sample MSE and correlation at the optimal bandwidth across 50 random train-test splits. For each kernel method in each split, we only keep the best out-of-sample results tuned over 10 kernel bandwidths and 32 numbers of PCs. Table 1 shows that smoothed connectomes produced by RDK and RMKs demonstrate superior predictive performance compared to SHK in predicting reading recognition scores. They also have better or competitive performance compared to SHK in predicting crystallized cognition composite, cognitive function composite (total composite) and picture vocabulary scores.

**Table 1.**
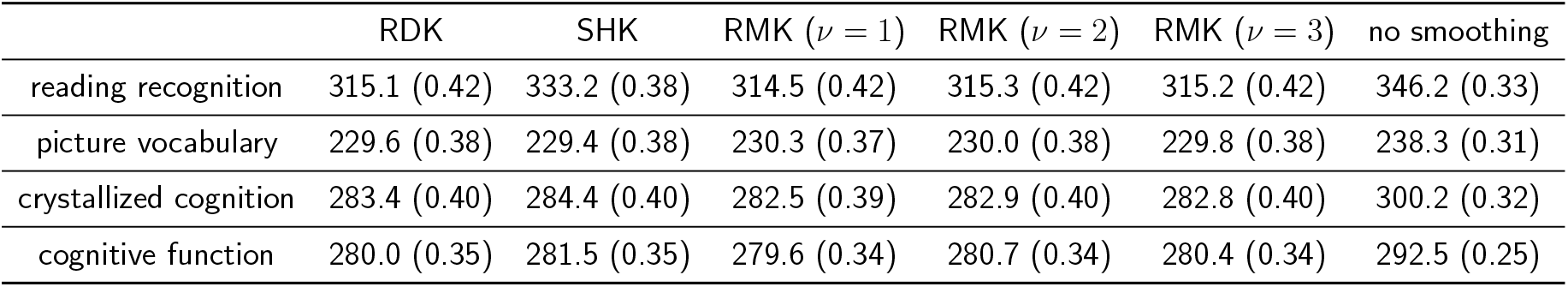
Cognitive trait prediction results of different kernel smoothing methods. Each cell shows the average out-of-sample MSE and correlation (in parentheses) between the observed and predicted outcomes over 50 random train-test splits, where the kernel bandwidth and the number of PCs are set at the optimal values.

| | RDK | SHK | RMK ($\nu = 1$) | RMK ($\nu = 2$) | RMK ($\nu = 3$) | no smoothing |
| --- | --- | --- | --- | --- | --- | --- |
| reading recognition | 315.1 (0.42) | 333.2 (0.38) | 314.5 (0.42) | 315.3 (0.42) | 315.2 (0.42) | 346.2 (0.33) |
| picture vocabulary | 229.6 (0.38) | 229.4 (0.38) | 230.3 (0.37) | 230.0 (0.38) | 229.8 (0.38) | 238.3 (0.31) |
| crystallized cognition | 283.4 (0.40) | 284.4 (0.40) | 282.5 (0.39) | 282.9 (0.40) | 282.8 (0.40) | 300.2 (0.32) |
| cognitive function | 280.0 (0.35) | 281.5 (0.35) | 279.6 (0.34) | 280.7 (0.34) | 280.4 (0.34) | 292.5 (0.25) |

### 3.5 Local Inference Detection Between Groups with High and Low Reading Ability

Figure 10 indicates that kernel-smoothed connectomes at appropriate bandwidths have strong predictive power for reading recognition scores. Therefore, we narrow our focus in this section to analyze the core cognition trait - *reading recognition*. Our goal is to identify the regions on the cortical surface Ω, where continuous connectivity differs between groups with high and low reading ability. The two groups were created by selecting the top 20% and bottom 20% of the 318 ABCD subjects based on their reading recognition scores from the baseline year, resulting in 62 subjects in each group. Locally supported basis functions *ξ*_*k*_’s (as described in (12) with *K* = 100) were estimated using the full set of 318 ABCD subjects to improve estimation accuracy, and the local inference procedure outlined in Section 2.6 was then applied.

The coefficient associated with the basis function *ξ*_1_ was found to be significantly different between the high and low reading ability groups when using the RDK, but no significant coefficients were observed under the SHK. Figure 11(a) shows the support set of *ξ*_1_ located on the right hemisphere of the white surface when using RDK. For each kernel, the KDE bandwidth was set to minimize the average out-of-sample MSE for reading recognition prediction, as illustrated in Figure 10.

**Figure 11.**
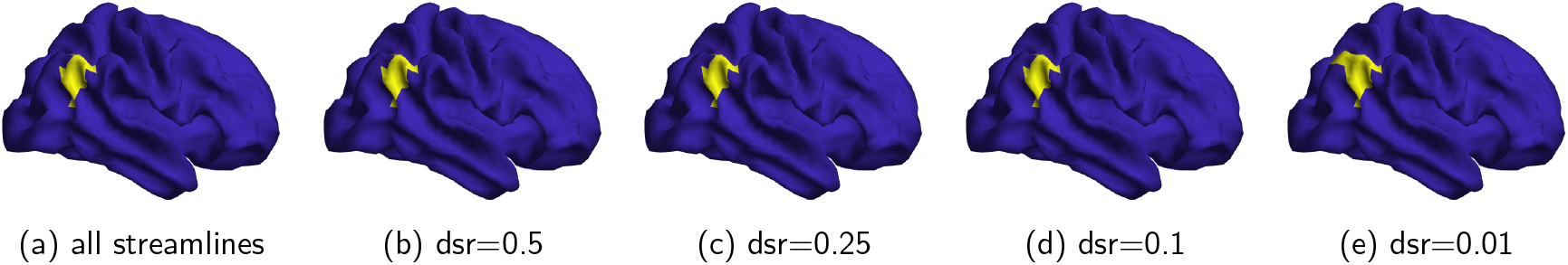
The support set (yellow) of the basis function *ξ*_1_ found to be significantly associated with connectivity differences in the high vs low reading ability groups using the Riemannian diffusion kernel over different streamline downsampling rates (dsr).

### 3.6 Impact of Downsampling Streamlines on Cognitive Trait Prediction and Local Inference

We first investigate how trait prediction performance alters when using structural density functions estimated from downsampled streamlines. For each downsampling rate in {50%, 25%, 10%, 5%, 1%}, we randomly sampled the total number of streamlines multiplied by the downsampling rate without replacement from the original tractography for each subject. We then estimated a high-resolution connectivity matrix (15) using the downsampled streamlines for each subject, conducted PCA, and entered the principal components in ridge regression for predicting each cognitive trait. Additionally, we report the results using the full set of extracted streamlines, corresponding to a downsampling rate of 100%, for comparison.

Figure 12 displays the average out-of-sample MSE for varying downsampling rates and bandwidths using RDK and SHK, respectively (see Supplementary Figure S7 for the correlation results). It is observed that, across all bandwidths and the four cognitive traits, the average predictive performance remains relatively consistent as the downsampling rate declines from 100% to 10% for both RDK and SHK. However, as the downsampling rate decreases to 5% or 1%, there is a noticeable increase in the average prediction errors. For instance, under the RDK method, the prediction errors for reading recognition scores increase significantly when the downsampling rate drops to 5% or lower. Similarly, for both RDK and SHK, the prediction errors for picture vocabulary scores become markedly worse at a downsampling rate of 1%.

**Figure 12.**
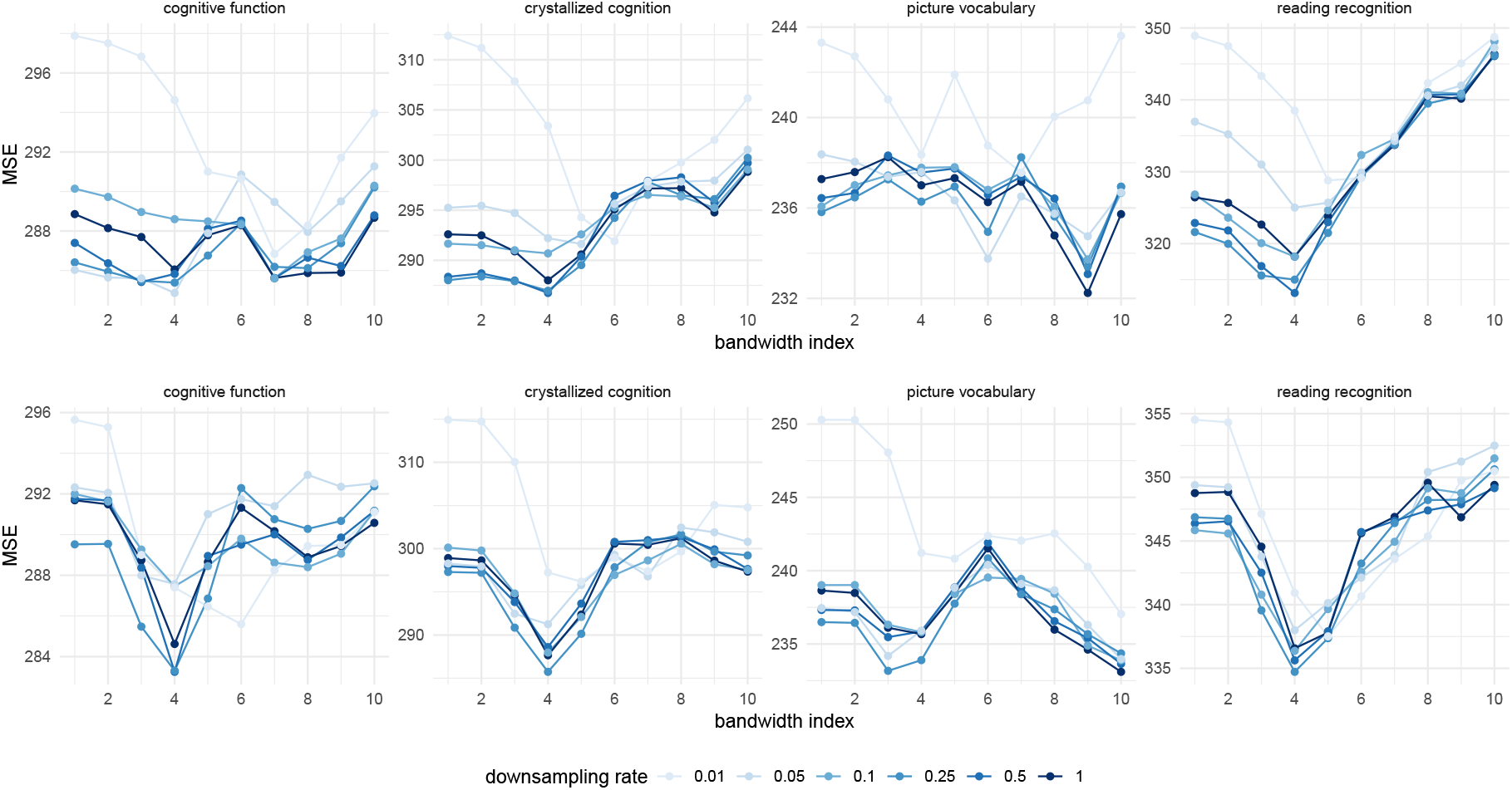
Average out-of-sample MSE between the observed and predicted cognitive measures, across various bandwidths at different downsampling rates. The first row displays results using the Riemannian diffusion kernel (RDK), while the second row shows results for the spherical heat kernel (SHK).

Table 2 presents the average out-of-sample MSE and correlation at the optimal bandwidth across different downsampling rates for both the RDK and SHK methods. We observe that the predictive performance for all four cognitive traits remains relatively stable when downsampling to 10% of the original tractography using either kernel. However, there is a significant decline in performance when the downsampling rate falls below 5%.

**Table 2.** Cognitive trait prediction results for different downsampling rates under the Riemannian diffusion kernel (RDK) and the spherical heat kernel (SHK) respectively. Each cell shows the average out-of-sample MSE and correlation (in parentheses) between the observed and predicted outcomes over 50 random train-test splits, where the kernel bandwidth and the number of PCs are set at the optimal values.

| downsampling rate | 100% | 50% | 25% | 10% | 5% | 1% |
| --- | --- | --- | --- | --- | --- | --- |
| RDK |  |  |  |  |  |  |
| reading recognition | 315.1 (0.42) | 310.1 (0.43) | 311.2 (0.43) | 314.9 (0.42) | 319.7 (0.41) | 325.1 (0.40) |
| picture vocabulary | 229.6 (0.38) | 230.1 (0.37) | 229.7 (0.38) | 230.7 (0.37) | 229.1 (0.37) | 234.3 (0.37) |
| crystallized cognition | 283.4 (0.40) | 281.7 (0.39) | 282.1 (0.40) | 285.4 (0.39) | 286.5 (0.39) | 288.7 (0.39) |
| cognitive function | 280.0 (0.35) | 279.5 (0.34) | 279.2 (0.35) | 281.8 (0.34) | 279.9 (0.35) | 284.4 (0.33) |
| SHK |  |  |  |  |  |  |
| reading recognition | 333.2 (0.38) | 332.6 (0.38) | 331.3 (0.38) | 333.6 (0.38) | 334.6 (0.37) | 334.2 (0.38) |
| picture vocabulary | 229.4 (0.38) | 229.8 (0.39) | 229.1 (0.39) | 230.1 (0.38) | 228.9 (0.38) | 235.5 (0.36) |
| crystallized cognition | 284.4 (0.40) | 285.1 (0.39) | 282.6 (0.40) | 284.7 (0.39) | 286.3 (0.40) | 291.5 (0.40) |
| cognitive function | 281.5 (0.35) | 280.7 (0.35) | 280.2 (0.35) | 282.8 (0.35) | 284.4 (0.35) | 280.3 (0.36) |

To evaluate the consistency of the identified regions from the local inference test across different downsampling rates, we focus once again on the local inference detection between groups with high and low reading ability. The test procedure outlined in Section 3.5 was repeated for each downsampling rate in the set {50%, 25%, 10%, 5%, 1%} using both RDK and SHK. In each case, the kernel bandwidth was set to minimize the average out-of-sample MSE for reading recognition scores, as illustrated in Figures 12.

For RDK-smoothed densities, the support regions of significant basis functions remain unchanged when the downsampling rate decreases from 100% to as low as 10%, as illustrated in Figures 11(a) through (d). At a downsampling rate of 5%, no significant basis components are identified. However, when the downsampling rate is further reduced to 1%, one basis function becomes significant, albeit with a slightly altered support region, as depicted in Figure 11(e).

For SHK-smoothed densities, no significant basis components are identified when using all streamlines down to a downsampling rate of 5%. However, at a downsampling rate of 1%, one basis function emerges as significant, sharing an identical support region with the RDK method at the same downsampling rate, as shown in Figure 11(e). downsampling rate 100% 50% 25% 10% 5% 1%

## 4 Discussion

In this paper, we propose an efficient method for constructing continuous structural connectivity directly on the white surface of the brain. This approach leverages recently developed techniques to compute the Riemannian diffusion kernel (RDK) on compact Riemannian manifolds (Borovitskiy et al., 2020). Our method effectively integrates the geometry of the cortical surface and demonstrates computational efficiency and improved performance in cognitive trait prediction. The following discussion synthesizes our findings to provide practical guidelines for researchers, contextualize the methodological advantages of our approach, and interpret its neuroscientific implications.

### 4.1 Kernel Bandwidth Selection for ABCD Data

Our evaluation of four distinct bandwidth selection criteria reveals a critical trade-off and offers clear guidance for future studies. The reliability measures dICC and *µ*_*intra*_ improve consistently as the kernel bandwidth increases, leading to the selection of excessively large kernel bandwidths, and resulting in oversmoothing of the data. Conversely, the identifiability measure proposed by Mansour L et al. (2022) shows a decrease with increasing kernel bandwidth, resulting in the selection of extremely small bandwidths. Given these findings, we suggest that trait prediction might offer a more reliable criterion for determining the optimal kernel bandwidth. For example, when predicting reading recognition scores, Figure 10 shows that both RDK and SHK achieve their best average out-of-sample performance at the fourth bandwidth level. These bandwidths seem to strike a balance between smoothing and signal integrity preservation, as illustrated in Figure 8.

However, it is important to note that the identified optimal bandwidths for RDK or SHK are not universally applicable. The most appropriate bandwidth can vary significantly depending on the specific neuroimaging dataset and the particular downstream tasks being performed. Therefore, tailored bandwidth selection is essential for different applications to ensure optimal results.

### 4.2 Advantages of the Riemannian Diffusion Kernel

Our comprehensive comparative analysis of RDK and SHK highlights several advantages of using RDK. First, RDK proves to be more time-efficient in computing high-resolution kernel matrices and kernel-smoothed connectivity matrices on a fine mesh. This efficiency can significantly reduce processing time when constructing continuous structural connectivity, particularly in large-scale neuroimaging studies.

Second, simulation studies indicate that RDK offers superior potential for more accurate estimation of connectivity density with fewer extracted streamlines, particularly when the endpoint pairs follow a mixture of “normal distributions” on the white surface. Practical considerations, along with the similar predictive performance of RDK and the Riemannian Matérn kernels (RMKs), as shown in Figure 10 and Table 1, suggest that streamline endpoints are more likely to follow a mixture of “normal distributions” on the white surface rather than on a sphere.

Third, regarding predictive performance, RDK-smoothed continuous connectivity significantly outperforms SHK-smoothed connectivity in predicting reading recognition scores. Additionally, RDK demonstrates competitive performance in predicting picture vocabulary scores compared to SHK. As a result, RDK achieves better overall performance in predicting crystallized cognition composite and cognitive function (total composite), as evidenced by Table 1.

Fourth, only RDK-smoothed densities produce a significant basis function in the local inference test between groups with high and low reading abilities. This finding allows for the identification of regions on the white surface where continuous connectivity differs significantly between the two groups. In contrast, SHK does not yield any significant components in this test.

### 4.3 The Number of Streamlines Required for Reliable Continuous Connectivity Estimation

Our findings on streamline downsampling provide important practical guidance for optimizing tractography pipelines. In our investigation of the impact of downsampling streamlines on cognitive trait prediction and the local inference test, we found that estimating connectivity densities from downsampled streamlines at a rate of 10% generally does not compromise out-of-sample prediction errors for key cognitive traits or the results of the two-group local inference test. Given that the original number of streamlines extracted per subject averages approximately 5.2 *×* 10^5^, our findings suggest that it is feasible to extract as few as 50,000 streamlines in tractography without adversely affecting the results. This holds true whether using RDK or SHK for trait prediction tasks or the local inference test.

It is important to note that the results regarding the number of streamlines presented in this study may be influenced by the choice of tractography method employed. Variations in diffusion processing, including the local diffusion model and tracking algorithm, may yield different results.

### 4.4 Structural Connectivity Differences between Adolescents with High and Low Reading Ability

Beyond methodological advances, our framework successfully validated its real-world utility by identifying neuroanatomical substrates of reading ability. The local inference test, powered by the RDK-smoothed connectome, identified a specific region on the white surface of the right hemisphere, among which the probability densities of forming streamlines differ significantly between 9-10-year-old adolescents with high and low reading abilities. The identified region is located at the intersection of the temporal, parietal, and occipital lobes of the right hemisphere, which is consistent with the existing understanding that several brain regions must work together for successful reading (Buchsbaum et al., 2005). Specifically, the top, back part of the temporal lobe (the parietal-temporal cortex) is responsible for phoneme analysis and phoneme-grapheme association, which drives the phonological processing system. The temporal-occipital cortex is where visual word forms and this area of the brain drives the orthographic processing system. In particular, the region we identified is near the planum temporale, an area at the junction of the parietal, occipital, and temporal lobes. This area is known to facilitate the rapid coordination of phonological and orthographic processing systems when we read text (Woollams et al., 2018).

### 4.5 Future Work

Our proposed method for constructing continuous structural connectivity provides interesting opportunities for future research, particularly in transitioning from network-based approaches to function-based methodologies. Potential directions for further exploration include: (i) establishing a robust and more generic criterion for kernel bandwidth selection. Rather than relying on a single task, one could adopt a multi-task learning framework to identify a bandwidth that yields stable and accurate predictions across a diverse set of cognitive and behavioral traits. This approach would ensure the selected bandwidth is not overfit to a specific outcome and instead captures neurobiological features with broad predictive utility; (ii) developing functional regression models that associate cognitive traits with continuous connectivity objects, with an additional focus on identifying predictive sparse regions on the white surface of the brain; (iii) developing hypothesis testing procedures to detect localized regions on the white surface where connectivity patterns vary between two time points during adolescence; and (iv) developing longitudinal functional regression methods to analyze the evolution of connectivity patterns over time, both at the population level and the individual level.

## Data and Code Availability

The data and code of this study are available from the corresponding author, upon reasonable request.

## Author Contributions

L.W.: Formal analysis, Visualization, Writing - original draft; D.L.: Conceptualization, Methodology, Writing - review & editing; Z.Z.: Conceptualization, Data curation, Supervision, Writing - review & editing.

## Supplementary Material

Supplementary material for this article is available in a separate file.

