## Supplemental Material for "Riemannian Diffusion Kernel-smoothed Continuous Structural Connectivity On Cortical Surface"

#### Supplementary Materials for "Riemannian Diffusion Kernel-smoothed Continuous Structural Connectivity On Cortical Surface"

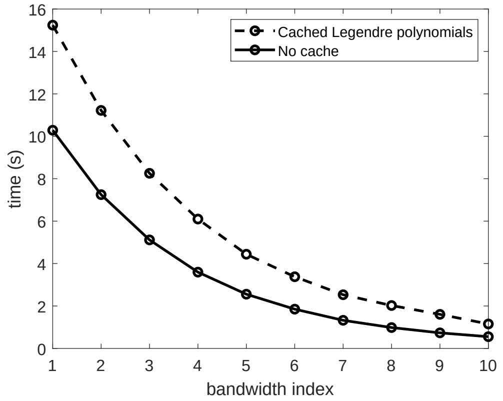


Figure S1: Average CPU time over 10 replications for computing the spherical heat kernel (SHK) matrices $\boldsymbol{K}_L$ and $\boldsymbol{K}_R$ in Equation (9) across 10 pre-specified bandwidths. 'Cached Legendre polynomials' refers to the scenario where the Legendre polynomial values for pairwise vertices are computed in advance and stored. 'No cache' indicates the scenario where the Legendre polynomial values are recomputed for each bandwidth without caching.

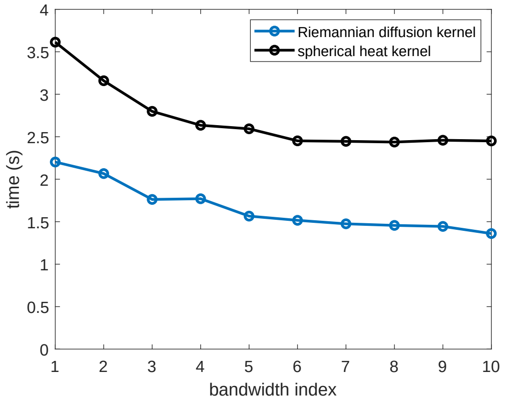


Figure S2: Average CPU time for computing the kernel-smoothed connectivity matrix $\boldsymbol{P}$ in Equation (15) for each individual scan using two kernels under each respective bandwidth.

### S1 Simulation Study: Evaluating Consistency in Kernel Density Estimation

In this section, we assess the performance of KDE using different numbers of streamline endpoints generated from a known distribution. To do this, we consider a random endpoint pair $(\boldsymbol{s}_1, \boldsymbol{s}_2)$ that follows a two-bundle mixture distribution. With a probability of 0.5, $\boldsymbol{s}_1$ and $\boldsymbol{s}_2$ are centered around $\boldsymbol{c}_1$ and $\boldsymbol{c}_2$, respectively, and with the same probability, they are centered around $\boldsymbol{c}_3$ and $\boldsymbol{c}_4$ on the cortical surface. The endpoints surrounding each center $\boldsymbol{c}_j$ are modeled by a "normal distribution" on the white surface $\Omega$, where the endpoint density is proportional to the RDK $k_{\kappa^*}(\boldsymbol{c}_j, \cdot)$ with $\kappa^* = 6$ for $j = 1, 2, 3, 4$, as illustrated in Figure S3. Consequently, the true density function of an endpoint pair on $\Omega \times \Omega$ is given by:

$$\begin{aligned} p^*(\boldsymbol{x}, \boldsymbol{y}) \propto\ & 0.5 \left[k_{\kappa^*}(\boldsymbol{x}, \boldsymbol{c}_1) k_{\kappa^*}(\boldsymbol{y}, \boldsymbol{c}_2) + k_{\kappa^*}(\boldsymbol{y}, \boldsymbol{c}_1) k_{\kappa^*}(\boldsymbol{x}, \boldsymbol{c}_2)\right] + \\ & 0.5 \left[k_{\kappa^*}(\boldsymbol{x}, \boldsymbol{c}_3) k_{\kappa^*}(\boldsymbol{y}, \boldsymbol{c}_4) + k_{\kappa^*}(\boldsymbol{y}, \boldsymbol{c}_3) k_{\kappa^*}(\boldsymbol{x}, \boldsymbol{c}_4)\right]. \end{aligned} \tag{S1}$$

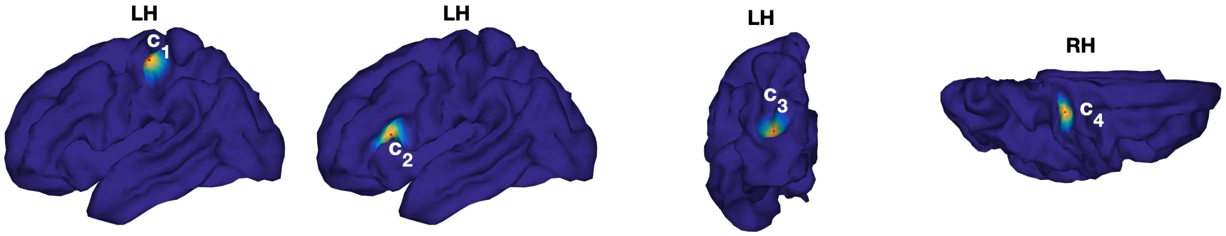


Figure S3: The Riemannian diffusion kernel (RDK) $k_\kappa(\boldsymbol{c}_j, \cdot)$ defined on the white surface with $\kappa = 6$ for $j = 1, 2, 3, 4$. Each center $\boldsymbol{c}_j$ is marked with a red dot.

Upon generating $N$ streamline endpoint pairs from (S1), we perform kernel density estimation and compute the error between the estimated density $\hat{p}(\boldsymbol{x}, \boldsymbol{y})$ (see (1)) and the true density $p^*(\boldsymbol{x}, \boldsymbol{y})$. The total estimated error over $\Omega \times \Omega$ is measured by the $L_2$ norm of $(\hat{p} - p^*)$, calculated as:

$$\begin{aligned} ||\hat{p} - p^*||_2 &= \left( \int_\Omega \int_\Omega [\hat{p}(\boldsymbol{x}, \boldsymbol{y}) - p^*(\boldsymbol{x}, \boldsymbol{y})]^2 d\boldsymbol{x} d\boldsymbol{y} \right)^{1/2} \\ &\approx \left\| \hat{\boldsymbol{P}} - \boldsymbol{P}^* \right\|_F \end{aligned} \tag{S2}$$

where $\hat{\boldsymbol{P}}$ and $\boldsymbol{P}^*$ are the discretized connectivity matrices of $\hat{p}$ and $p^*$ defined on a fine mesh of $\Omega \times \Omega$, and $\| \cdot \|_F$ is the Frobenius norm. The consistency of each kernel method can be assessed by checking whether the estimated error (S2) decreases consistently as the number of observed streamlines $N$ increases.

We vary the sample size $N$ over the discrete values $\{10, 50, 100, 200, 300, 400, 500\}$. For both the RDK and the Riemannian Matérn kernels (RMKs), the bandwidth parameter $\kappa$ is tuned among $\{1, 2, 3, 4, 5, 6, 7, 8, 9, 10\}$. To achieve comparable impact radii on a sphere, the bandwidth parameter $\sigma$ for the SHK is tuned over the set $\{0.0003, 0.001, 0.002, 0.003, 0.004, 0.006, 0.008, 0.01, 0.015, 0.02\}$.

For each sample size, we randomly generate 50 datasets of streamline endpoint pairs according to the true probability density function (S1). The left plot of Figure S4 displays the average errors of the kernel density estimation in relation to the sample size $N$ for each kernel method. The results reveal that the estimation error consistently decreases with increasing sample sizes for each kernel, confirming their consistency. For this specific mixture distribution, where streamlines form two bundles, RDK achieves the lowest average estimation error when $N < 200$ or each bundle contains fewer than 100 streamlines. For RMKs, the estimation errors decline as the parameter $\nu$ increases at each sample size, which is expected since endpoints near each local center follows a "normal distribution" on the white surface, and RMK converges to RDK as $\nu \to \infty$. When using the SHK, the density estimates yield the highest average error with $N = 10$ compared to the other kernels. This larger error is likely due to the isotropic decay of SHK on the sphere, which doesn't align well with the white surface topology. However, as the sample size increases, SHK gradually outperforms RMK and RDK. This improvement may stem from the available analytic form of SHK, which allows it to approximate any density function more accurately, given a sufficiently large sample size and a suitable bandwidth.

The right plot of Figure S4 presents the average CPU time required to compute high-resolution kernel-smoothed connectivity matrices (15) across 10 bandwidths for each kernel and specified number of streamlines. As shown, RDK and RMKs exhibit significantly better time efficiency than SHK. This efficiency is attributed to the simple computational form of the discretized kernel matrices for RDK and RMKs, as discussed in Section 3.2, and their more efficient calculation of kernel-smoothed connectivity matrices, as noted in the same section.

To qualitatively evaluate the accuracy of density estimates, we randomly select one dataset across several sample sizes and display the estimated density, $\hat{p}(\boldsymbol{c}_3, \cdot)$, on the white surface for both RDK and SHK in Figure S5. Compared to the true density $p(\boldsymbol{c}_3, \cdot)$ shown in the leftmost plot of Figure 6, the results indicated that RDK achieves a reasonably accurate estimate with just 50 streamlines, while SHK requires 500 streamlines to attain the correct density estimate without distinct errors on the neighboring gyri around the high-density region at the sulcus.

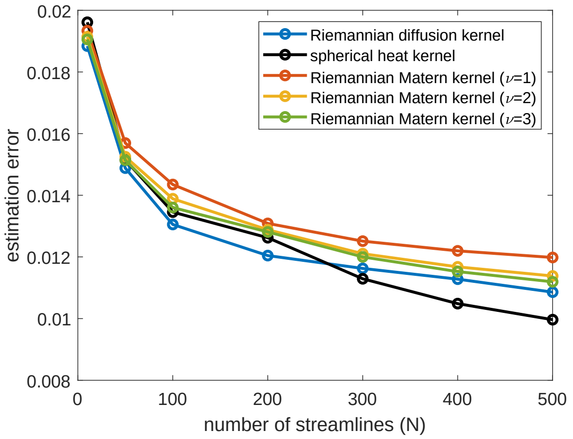


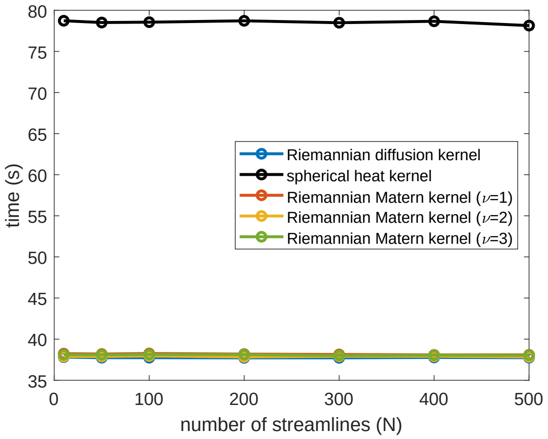


Figure S4: Left: Average estimated error over 50 replications versus the number of streamlines $N$ for different kernels. Right: Average CPU time to compute high-resolution kernel-smoothed connectivity matrices (15) over 10 bandwidths under each number of streamlines for different kernels.

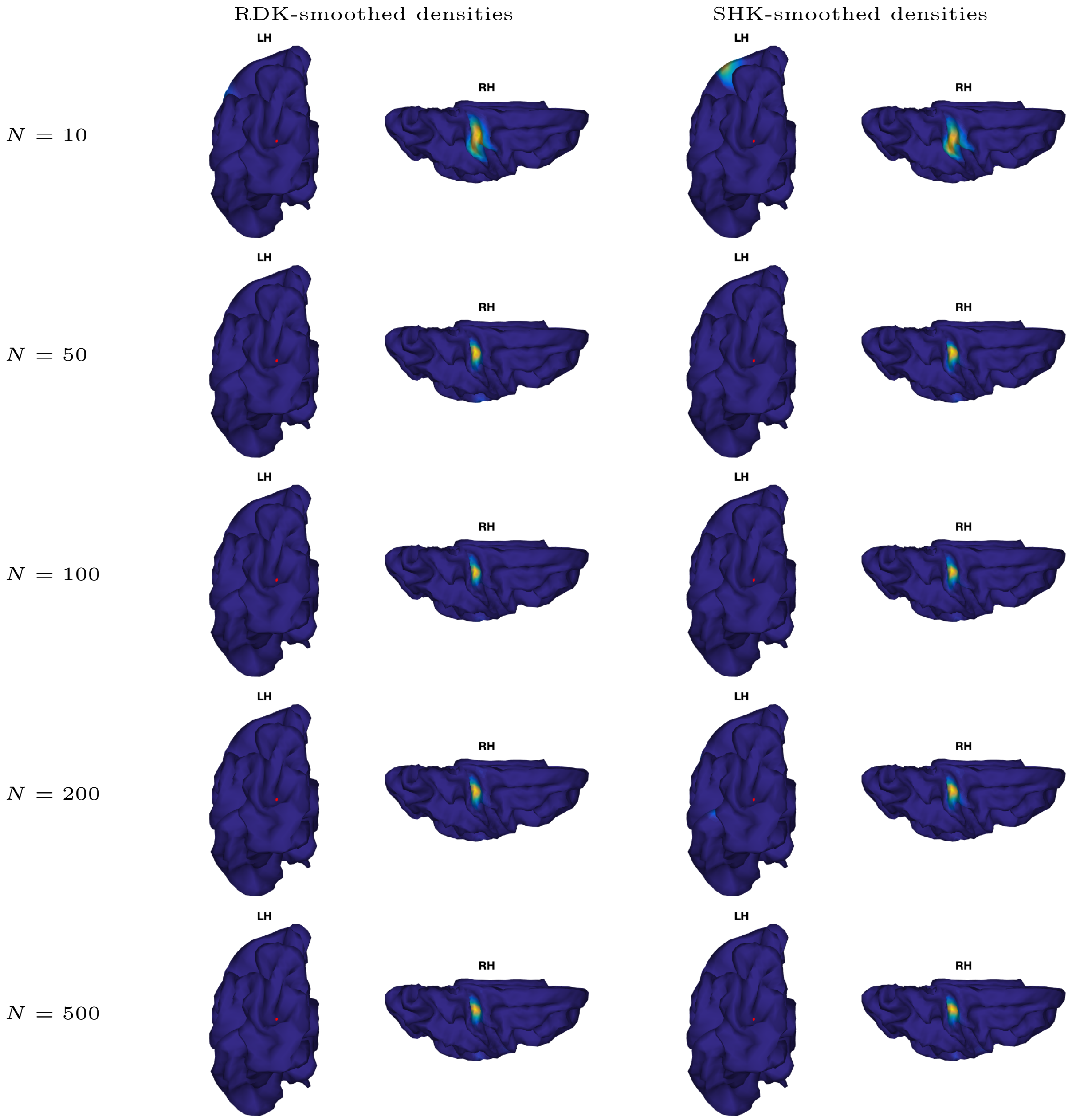


Figure S5: Estimated density $\hat{p}(\boldsymbol{c}_3, \cdot)$ using RDK (left) and SHK (right) for a dataset generated with different numbers of streamlines, $N$, in the simulation study. The point $\boldsymbol{c}_3$ is marked with a red dot and the color indicates the value of the function.

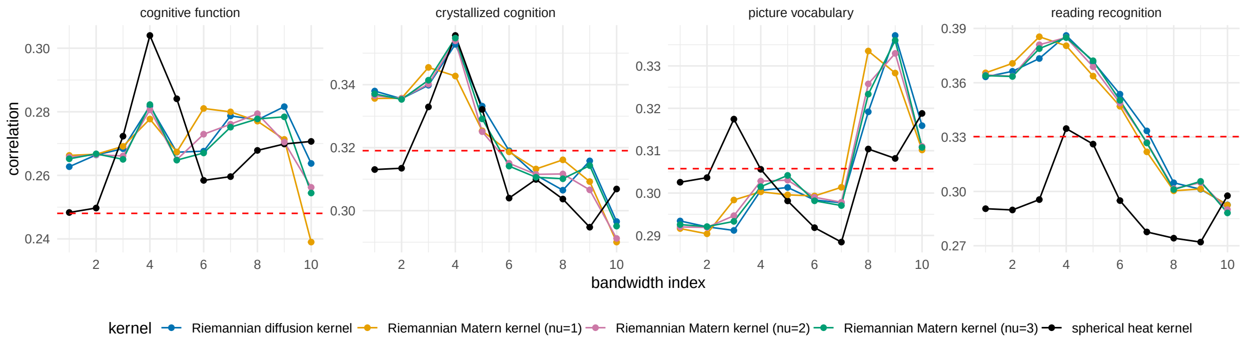


Figure S6: Average out-of-sample correlation between the observed and predicted cognitive measures, based on 50 random train-test splits, for five kernel methods across 10 bandwidths. The red dashed line represents the results for connectomes with no smoothing.

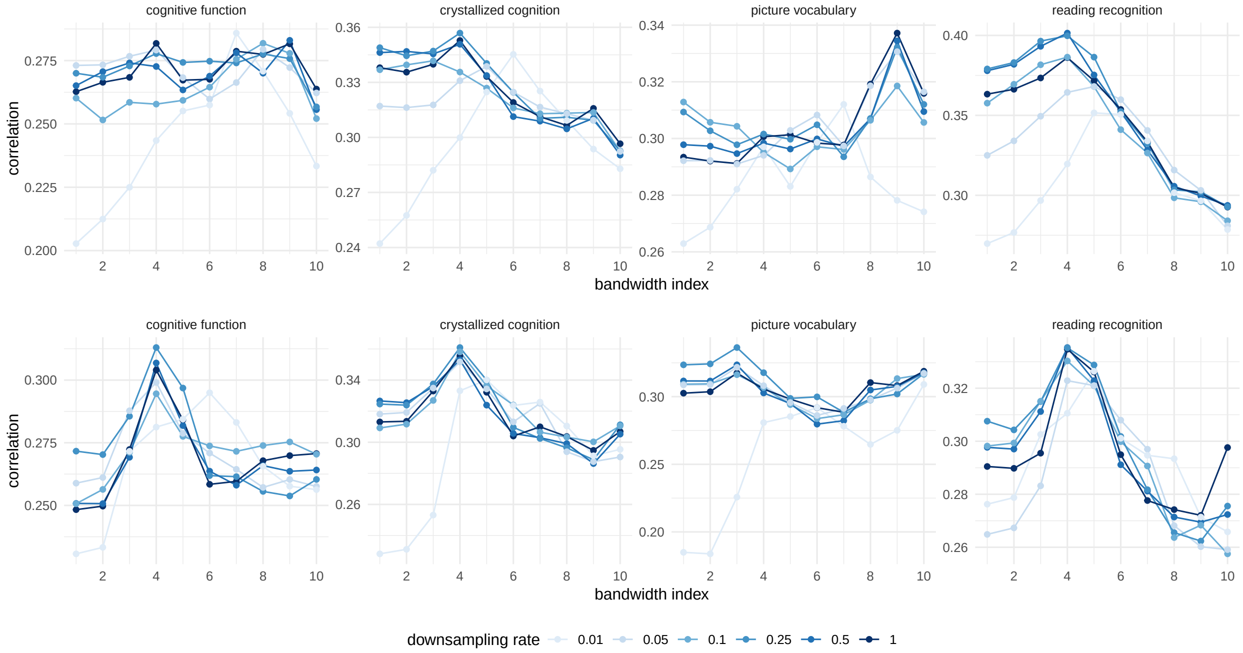


Figure S7: Average out-of-sample correlation between the observed and predicted cognitive measures, across various bandwidths at different downsampling rates. The first row displays results using the Riemannian diffusion kernel (RDK), while the second row shows results for the spherical heat kernel (SHK).
